# Computationally restoring the potency of a clinical antibody against SARS-CoV-2 Omicron subvariants

**DOI:** 10.1101/2022.10.21.513237

**Authors:** Thomas A. Desautels, Kathryn T. Arrildt, Adam T. Zemla, Edmond Y. Lau, Fangqiang Zhu, Dante Ricci, Stephanie Cronin, Seth J. Zost, Elad Binshtein, Suzanne M. Scheaffer, Bernadeta Dadonaite, Brenden K. Petersen, Taylor B. Engdahl, Elaine Chen, Laura S. Handal, Lynn Hall, John W. Goforth, Denis Vashchenko, Sam Nguyen, Dina R. Weilhammer, Jacky Kai-Yin Lo, Bonnee Rubinfeld, Edwin A. Saada, Tracy Weisenberger, Tek-Hyung Lee, Bradley Whitener, James B. Case, Alexander Ladd, Mary S. Silva, Rebecca M. Haluska, Emilia A. Grzesiak, Christopher G. Earnhart, Svetlana Hopkins, Thomas W. Bates, Larissa B. Thackray, Brent W. Segelke, Antonietta Maria Lillo, Shivshankar Sundaram, Jesse Bloom, Michael S. Diamond, James E. Crowe, Robert H. Carnahan, Daniel M. Faissol

## Abstract

The COVID-19 pandemic underscored the promise of monoclonal antibody-based prophylactic and therapeutic drugs^1–3^, but also revealed how quickly viral escape can curtail effective options^4, 5^. With the emergence of the SARS-CoV-2 Omicron variant in late 2021, many clinically used antibody drug products lost potency, including Evusheld^TM^ and its constituent, cilgavimab^4, 6^. Cilgavimab, like its progenitor COV2-2130, is a class 3 antibody that is compatible with other antibodies in combination^4^ and is challenging to replace with existing approaches. Rapidly modifying such high-value antibodies with a known clinical profile to restore efficacy against emerging variants is a compelling mitigation strategy. We sought to redesign COV2-2130 to rescue in vivo efficacy against Omicron BA.1 and BA.1.1 strains while maintaining efficacy against the contemporaneously dominant Delta variant. Here we show that our computationally redesigned antibody, 2130-1-0114-112, achieves this objective, simultaneously increases neutralization potency against Delta and many variants of concern that subsequently emerged, and provides protection *in vivo* against the strains tested, WA1/2020, BA.1.1, and BA.5. Deep mutational scanning of tens of thousands pseudovirus variants reveals 2130-1-0114-112 improves broad potency without incurring additional escape liabilities. Our results suggest that computational approaches can optimize an antibody to target multiple escape variants, while simultaneously enriching potency. Because our approach is computationally driven, not requiring experimental iterations or pre-existing binding data, it could enable rapid response strategies to address escape variants or pre-emptively mitigate escape vulnerabilities.

## Introduction

The COVID-19 pandemic has underscored the promise of monoclonal antibody-based drugs as prophylactic and therapeutic treatment options for infectious disease. Multiple monoclonal antibody drug products were developed and authorized for emergency use by the US FDA that demonstrated efficacy in preventing COVID-19^1^, reducing death and hospitalization rates^2^ or reducing viral load^3^.

Despite these efforts, SARS-CoV-2 variant Omicron BA.1 escaped many monoclonal antibody and antibody combination drug products deployed under emergency use authorization by the FDA^6, 7^. First reported in November 2021, BA.1 outcompeted all other VOCs worldwide within weeks^8^. BA.1 contains over 50 substitutions, including 15 in the spike protein receptor binding domain (RBD), the primary target for therapeutic and prophylactic antibodies. These substitutions reduce or eliminate the neutralization capacity of many authorized prophylactic and therapeutic antibodies^4, 5, 7^.

In particular, the antibody combination Evusheld^TM^–the only antibody drug approved for pre-exposure prophylaxis in immunocompromised patients for whom vaccination is not always protective^1^ –was impacted by the emergence of the Omicron variants. Evusheld combines tixagevimab plus cilgavimab, which are composed of the progenitor monoclonal antibodies COV2-2196 and COV2-2130, respectively. The two-antibody cocktail exhibits an approximately 10- to 100-fold reduction in neutralizing potency against Omicron BA.1 compared to wild-type SARS-CoV-2^4, 9^. COV2-2130 suffers an approximately 1,000-fold loss in neutralization potency against Omicron BA.1.1 compared to strains circulating earlier in the pandemic^7, 10, 11^.

COV2-2130 is a class 3 RBD-targeting antibody that blocks the RBD-ACE2 interaction without competing with antibodies targeting the class 1 site on RBD. Thus, class 1 and class 3 antibodies can be combined or co-administered for simultaneous binding and synergistic neutralization^12^. While antibodies that target the class 3 site of RBD have clear utility for use in therapeutic antibody combinations, the emergence of Omicron BA.1 and BA.1.1 reduced or abrogated the binding and neutralization of many antibodies currently available ^4^. Furthermore, potently neutralizing antibodies targeting class 3 sites on RBD are less frequently identified^12^, suggesting that they are more difficult to replace with existing approaches.

Computational re-design of a monoclonal antibody is a promising strategy to recover efficacy against escape variants. Its value is further enhanced in the case of an antibody that has demonstrated efficacy and safety in clinical trials and is known to be compatible and synergistic with other clinically used monoclonal antibodies as part of a combination antibody drug product, such as COV2-2130^12^. To this end, we sought to optimize COV2-2130 to restore potent neutralization of SARS-COV-2 escape variants by introducing a small number of mutations in the paratope and computationally assessing improvement to binding affinity. We developed and used a computationally driven approach, called Generative Unconstrained Intelligent Drug Engineering (GUIDE), that combines high-performance computing, simulation, and machine learning to co-optimize binding affinity against multiple antigen targets, such as RBDs from several SARS-CoV-2 strains, along with other critical attributes such as thermostability. The computational platform operates in a “zero-shot” setting, i.e., designs are created without iteration through, or input from, wet laboratory experiments on proposed antibody candidates, relatives, or other derivatives of the parental antibody (e.g., single-point mutants). While more challenging, this zero-shot approach, if successful, can enable rapidly producing efficacious antibody candidates optimized for multiple target antigens in response to immediate needs presented by escape variants. We used our computational platform over a three-week period to repair the activity of COV2-2130 against Omicron variants.

## Computational design

Our computationally driven antibody design platform leverages simulation and machine learning to generate mutant antibody sequences that are co-optimized for multiple critical properties, without requiring experimental feedback or pre-existing binding data (**Fig. 1**). The platform comprises three phases: problem formulation, computational design and selection of mutant antibody candidates, and experimental validation of proposed candidates.

**Figure 1.**
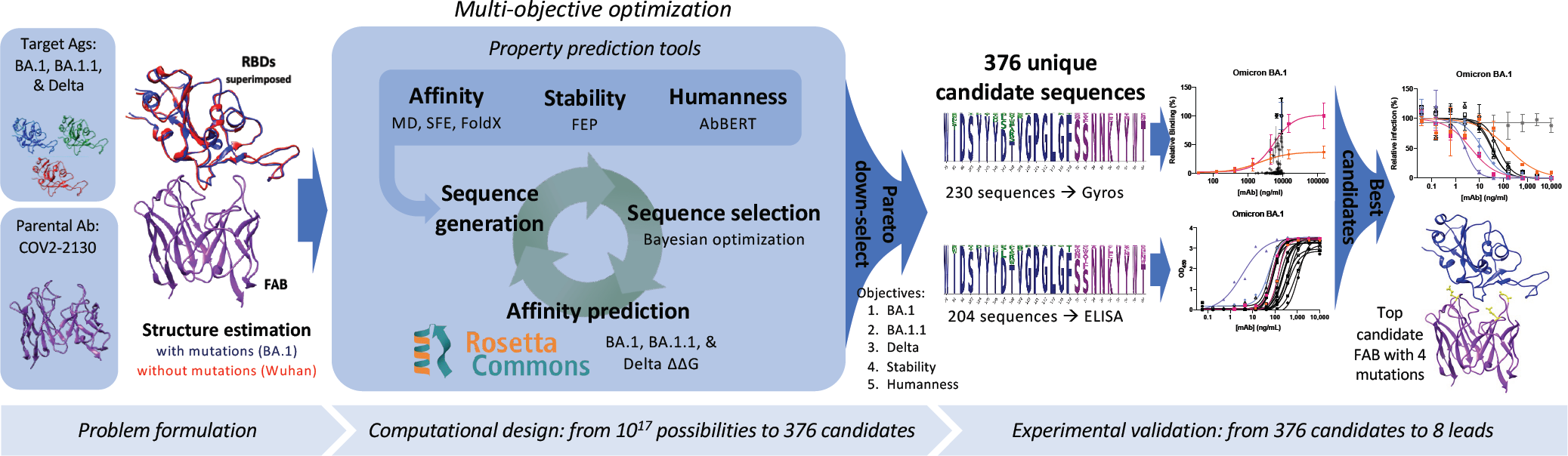
Overview of the GUIDE computationally driven drug engineering platform. Given a parental antibody and target antigens, co-structures are estimated experimentally and/or computationally (left). Within the main computational loop (center left), a sequence generator proposes multi-point mutant antibody candidates, and a Bayesian optimization agent selects which proposed sequences to evaluate via a set of affinity prediction tools. A subset of 376 computationally evaluated sequences based on Pareto optimality, mutational distance, and sequence diversity were experimentally evaluated for binding affinity by Gyros or ELISA (center right). The top sequences are then evaluated for neutralization of SARS-CoV-2 variants (right). See Methods for details.

We formulate a problem by identifying a parental antibody, a set of target antigens, and corresponding co-structures. In this case, we redesigned antibody COV2-2130^12^ for simultaneous binding improvements against Omicron BA.1 and BA.1.1 while maintaining binding to the Delta variant. We used co-structures that were both experimentally determined and computationally estimated, starting from co-structures including the wild-type antigen^13^. Since an experimental structure of the Omicron RBD was not available at the onset of our design process, we estimated the structure of the complex of the RBD with COV2-2130 using template-based structural modeling^14^. We incorporated experimentally determined Omicron RBD structures^15^ into the design process as they became available. We considered twenty-five paratope residues for mutation, primarily in or near the heavy (H) or light (L) chain complementarity determining regions (CDRs^16^) H2, H3, L1, and L2, and allowed up to nine amino acid substitutions per mutant sequence, resulting in a search space containing over 10^17^ possible mutant sequences.

Our computational design approach was implemented as a multi-objective optimization problem defined over this large space of mutations to COV2-2130 paratope residues. We considered five critical antibody properties: (1) binding affinity to Omicron BA.1 RBD, (2) binding affinity to BA.1.1 RBD, (3) binding affinity to Delta RBD, (4) thermostability, and (5) “humanness.” We expected restored affinity of a candidate antibody to each RBD variant to result in restored neutralization because the parental antibody, COV2-2130, competes with human angiotensin converting enzyme (ACE2) in SARS-CoV-2 spike binding^12^. Four complementary computational tools enable affinity prediction: atomistic potential of mean force molecular dynamics simulations (MD), Structural Fluctuation Estimation^17^ (SFE), Rosetta Flex^18^, and FoldX^19^. We estimated thermal stability using the Free Energy Perturbation (FEP) method^20^.

Humanness was quantified as the score under the AbBERT model^21^, a deep language model trained on a large database of human antibody sequences^22^. We include the AbBERT score as an objective based on the assumption that natural human immunoglobulin sequences are selected and expanded for being highly functional, and are therefore enriched for sequences that encode stable, soluble, non-autoreactive (i.e., more developable) antibodies. We used these tools to initialize a *sequence generator*, which proposes multi-residue mutations to the amino acid sequence of COV2-2130, biased toward residues that perform well across these critical properties. Within the optimization loop, this generator proposes batches of mutant antibody sequences. Next, we employed distributed software agents, each using Bayesian optimization or rules-based methods, to select a subset of promising candidate sequences to simulate in Rosetta Flex, yielding predicted binding affinities. Over the course of less than three weeks, we computationally evaluated more than 125,000 antibody candidates.

We calculated the Pareto optimal set^23^ based on the outputs of these tools, resulting in 3,809 sequences. Due to experimental capacity, we further down-selected from among the Pareto set based on mutational distance and sequence diversity. Our final selection of 376 antibody sequences was then advanced to experimental evaluation.

## Experimental evaluation

### Antibody and antigen production

We experimentally validated the 376 designed candidates. To leverage available resources at multiple experimental sites, we split candidates into two partially overlapping subsets, Sets 1 and 2. Set 1 consisted of 230 designs expressed as IgG in HEK-293 cells (ATUM), and Set 2 consisted of 204 designs expressed as IgG via a pVVC-mCisK_hG1 vector (Twist BioScience) in transiently transfected CHO cells. Omicron antigens were produced in Expi293F cells (ThermoFisher Scientific) and purified on HisTrap Excel columns (Cytiva).

In the following experiments, we selected antigens or viral strains to determine if we had achieved three goals. These goals consisted of (1) our primary design focus of improving binding affinity to and efficacy against BA.1 and BA.1.1; (2) maintaining efficacy against historical strains, for which design explicitly targeted Delta but experiments often substituted WA1/2020 D614G; and (3) determining whether our designs were robust to emerging VOCs.

### Computationally designed antibodies maintained favorable expression yields

Because *in silico* derivatization of antibody sequences can inadvertently compromise production yield, we measured concentrations of Set 1’s 230 COV2-2130-derived recombinant antibodies that were produced and compared these concentrations to that of the parental antibody produced in parallel. The purified concentrations of 73.9% of re-designed antibodies exceeded that of the parental COV2-2130 antibody (170/230 mAbs at >171.2 mg/L), reaching as high as 305 mg/L. Only approximately 10% of designed antibodies gave poor yields relative to the parental molecule (22/230 mAbs at <135 mg/L, i.e., <80% of parental antibody yield). Our designs thus yielded candidates for downstream characterization that retained fundamental production properties of the parental antibody.

### Computationally designed antibodies improved binding to Omicron subvariants and preserved thermostability

We screened all designed antibodies for binding to RBDs. For Set 1, we did so by a single-concentration immunoassay (Gyrolab xPlore) in the contexts of WA1/2020, Delta, BA.1, or BA.1.1 RBDs (see **Fig. ED1**). For Set 2, we used a multi-concentration immunoassay (ELISA; **Fig. ED2**) in the context of wild type, BA.1 or BA.1.1 RBDs. In the single-concentration immunoassay, this value was chosen as a single dilution factor causing most designed antibody samples to fall in the dynamic range of the positive control. In both cases, we compared the designed antibodies’ binding with a broadly cross-reactive control antibody S309^24^ and the parental COV2-2130 antibody. As intended, most antibody designs had altered binding profiles, indicating that the designed mutations were consequential. Approximately 11% of Set 1’s designs retained WA1/2020 antigen binding at the measured concentration; roughly 6% improved binding against BA.1, and 5% did so against BA.1.1. The corresponding numbers for Set 2 were 9% against WA1/2020 and 8% against BA.1. Following this initial screen, we down-selected both sets of antibody designs to those with improved binding to Omicron subvariants BA.1 and BA.1.1 for further characterization.

These down-selected antibodies were remanufactured at larger scale. We characterized the resulting IgG antibodies by immunoassay and thermal shift (melt temperature) assessments. In agreement with our screens, seven of the eight top-performing antibodies preserved comparable binding to WA1/2020 and Delta RBDs, improving over the parental COV2-2130 antibody with respect to their binding to Omicron BA.1 and BA.1.1 RBDs (**Fig. 2**). Furthermore, seven of the eight antibodies had melting temperatures and expression properties comparable to those of COV2-2130. One antibody, 2130-1-0114-111, had reduced melting temperature. (**Table EDT1**). Antibody 2130-1-0114-112 displayed best-in-class binding across all RBD variants and had no significant difference in thermal stability compared to the parental COV2-2130 antibody.

**Figure 2.**
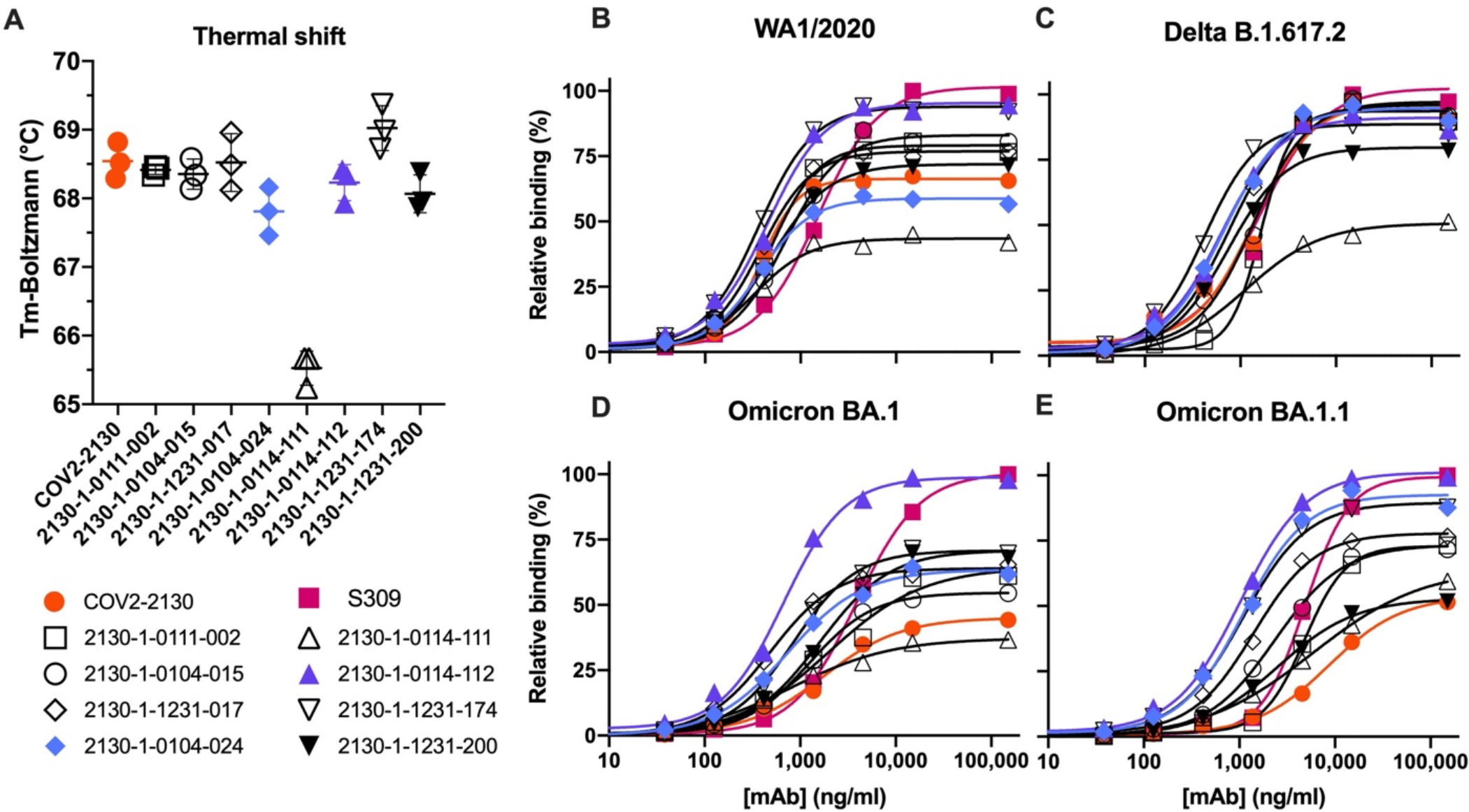
Computationally designed IgG antibodies improve Omicron binding and maintain parental thermostability and binding to historical strains. (A) The parental COV2-2130 (orange circles) and computationally designed antibodies (2130-1-0114-112, purple triangles; 2130-1-0104-024 blue diamonds; remainder in black) were assayed for thermal shift (n=3, technical replicates). Bars indicate the mean, and error bars indicate standard deviation. (B-E) The parental COV2-2130 antibody and computationally designed antibodies (represented by the same symbols as in A) and cross-reactive positive control antibody S309 (magenta squares) were analyzed for relative binding against four SARS-CoV-2 Spike-RBD variants in Gyrolab immunoassay: WA1/2020 (B), Delta (C), Omicron BA.1 (D) and Omicron BA.1.1 (E). Lines represent 4-parameter logistic regression model fit using GraphPad Prism to each titration, executed without technical replicates.

### Computationally designed antibodies restored neutralization to Omicron subvariants in pseudoviral neutralization assays

We performed pseudovirus neutralization assays to characterize the functional performance of five selected antibody designs (**Fig. 3**), comparing with parental COV2-2130, positive control S2K146^25^, and negative control DENV-2D22^26^. Our designs maintained neutralization activity against pseudoviruses displaying historical spike proteins (WA1/2020 D614G) and achieved neutralization of those with Omicron BA.1 spikes. The single best candidate design, 2130-1-0114-112, restored potent neutralization in the context of BA.1.1 and showed a two-order-of-magnitude improvement in IC50 versus parental COV2-2130 for BA.1 and BA.4. These pseudovirus neutralization test results showed that our designs neutralized BA.2 and BA.4 more potently than COV2-2130, despite the emergence of these VOCs after the conception of our designs. We additionally tested 2130-1-0114-112’s performance against BA.2.75, BA.4.6 (which contains an R346T mutation, among others), and an artificially constructed BA.2.75 + R346T, which matches the RBD sequence of BA.2.75.6 (**Fig. ED3**). 2130-1-0114-112 outperforms COV2-2130, including maintaining potent neutralization of BA.2.75 (IC50 of 2.6 ng/ml), which reduces COV2-2130’s potency by more than 30-fold. The R346T mutation leads to significant loss of potency for both parental COV2-2130 and 2130-1-0114-112, though 2130-1-0114-112 retains some measurable neutralization against BA.4.6 (IC50 of 1264 ng/ml) and BA.2.75 + R346T (IC50 of 673.8 ng/ml).

**Figure 3:**
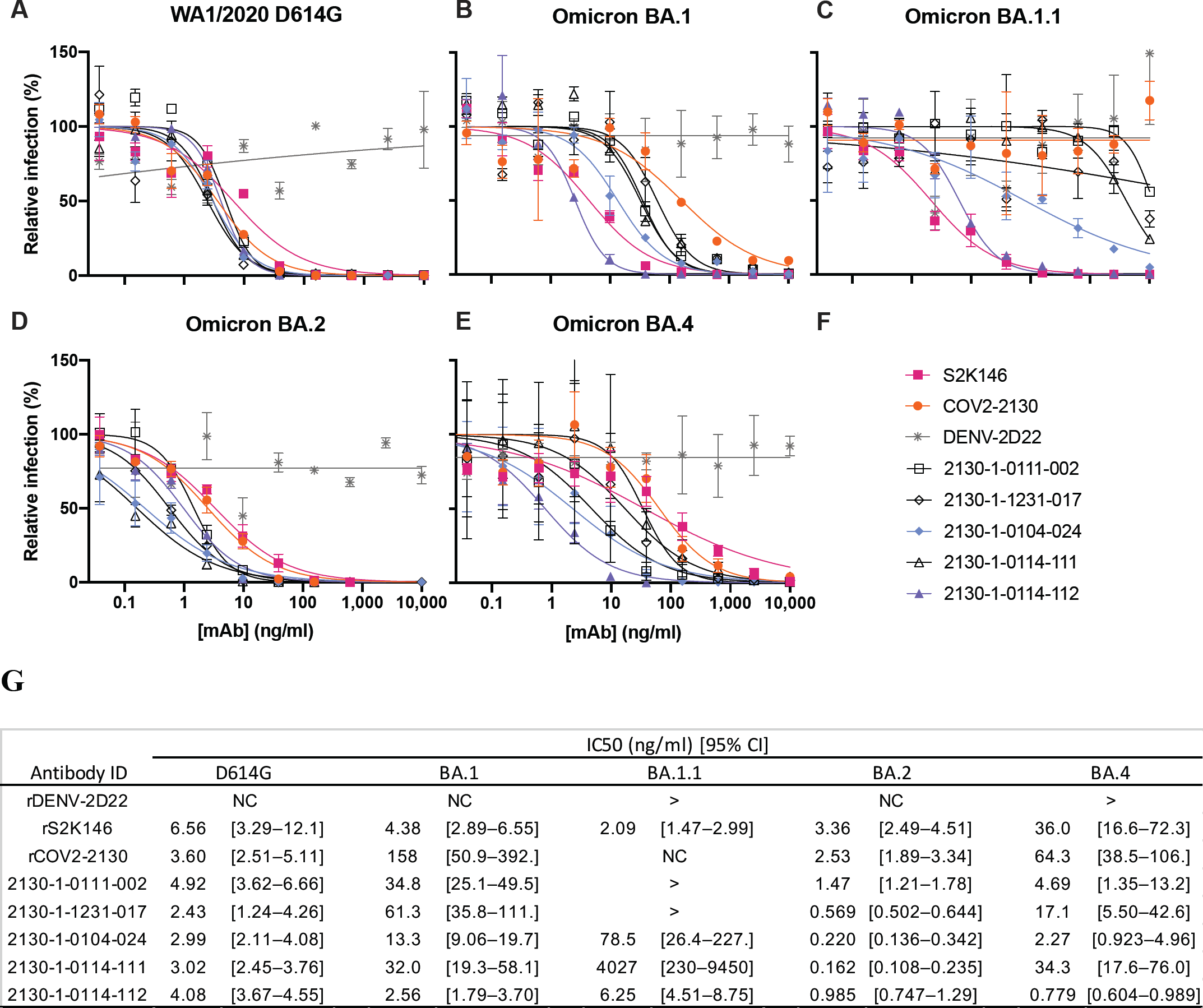
Designed antibodies improve pseudoviral neutralization over COV2-2130. The parental COV2-2130 antibody (orange circles), cross-reactive positive control antibody S2K146 (magenta squares), negative control antibody DENV-2D22 (gray x), and down-selected computationally designed antibodies were assayed by neutralization with lentiviruses pseudotyped with spike variants of WA1/2020 D614G (A), Omicron BA.1 (B), BA.1.1 (C), BA.2 (D), and BA.4 (E). (F) Symbols for each antibody are indicated in the legend. (G) IC50 values and 95% confidence intervals. “>” indicates a value > 10,000; NC indicates positive hill slope or failure to converge. Symbols indicate the mean and standard deviation of two technical replicates; curves are 4-parameter logistic regression models fit using GraphPad Prism.

### Top computationally designed antibody, 2130-1-0114-112, restores neutralization of Omicron subvariants in an authentic virus assay

We evaluated our best antibody, 2130-1-0114-112 (containing four mutations: GH112E, SL32A, SL33A, TL59E) for authentic virus neutralization performance against several strains of SARS-CoV-2 by a focus reduction neutralization test (FRNT) in Vero-TMPRSS2 cells (**Fig. 4**). The strains we used included several Omicron targets: WA1/2020 D614G, Delta (B.1.617.2), BA.1, BA.1.1, BA.2, BA.2.12.1, BA.4, BA.5, and BA.5.5. In all cases apart from Delta, 2130-1-0114-112 had an IC50 < 10 ng/ml. Compared to the parental COV2-2130, 2130-1-0114-112 restored potent neutralization activity against both BA.1 (8.08 ng/ml) and BA.1.1 (7.77 ng/ml), showed a more than 5-fold improvement in IC50 against BA.2 (2.4 ng/ml) and BA.2.12.1 (2.27 ng/ml), and conferred 50-fold or greater improvements in IC50 against BA.4 (3.16 ng/ml), BA.5 (3.51 ng/ml), and BA.5.5 (5.29 ng/ml). We also evaluated 2130-1-0114-112 and a less-mutated alternative design, 2130-1-0104-024 (containing two mutations: SL32W, TL59E), in plaque assays with Vero E6-TMPRSS2-T2A-ACE2 cells (**Fig. ED5**). IC50 values for 2130-1-0104-024 were 37.7 ng/ml, 75.94 ng/ml, and 781.7 ng/ml for Delta, BA.1, and BA.1.1 viruses, respectively.

**Figure 4:**
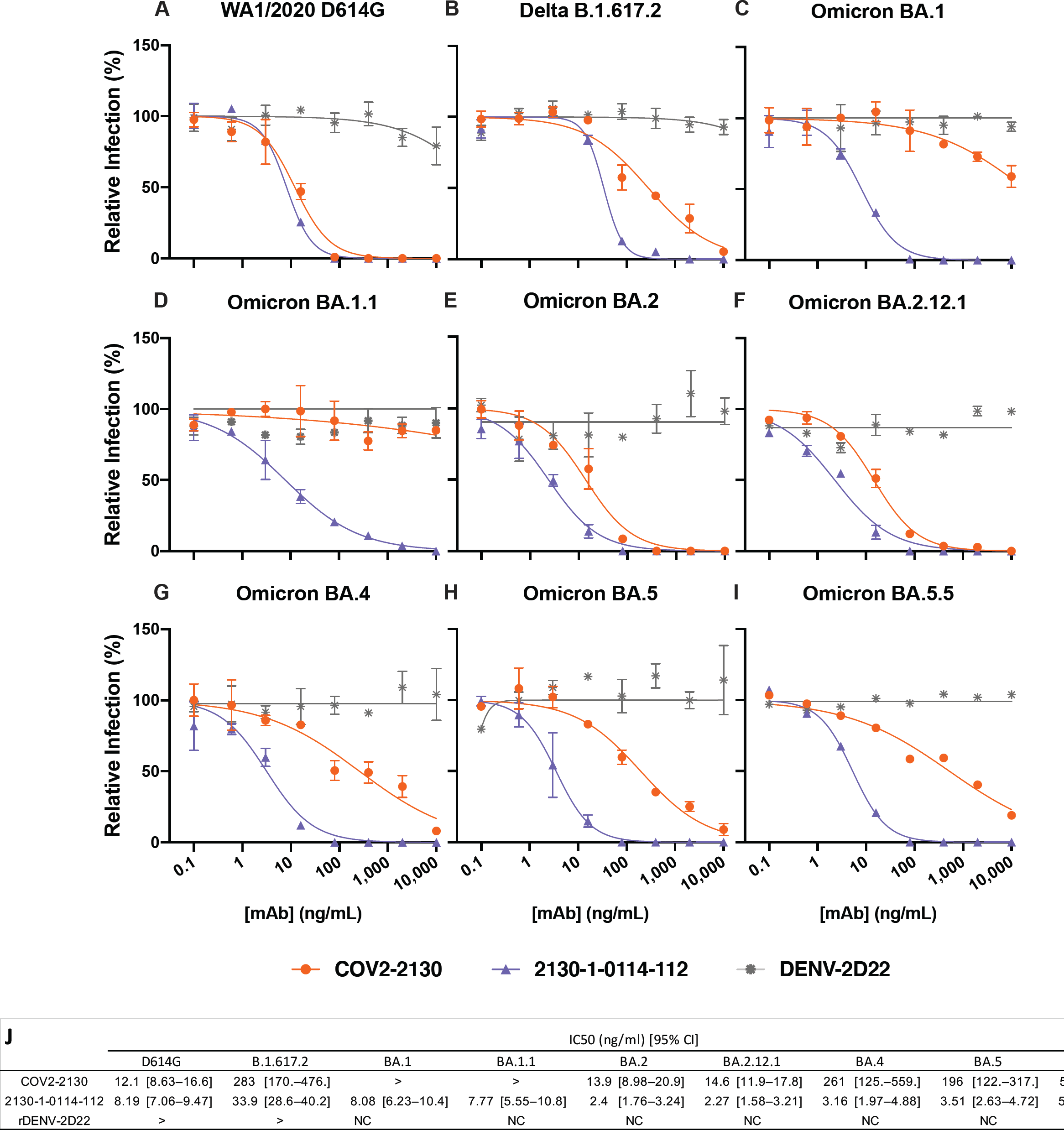
2130-1-0114-112 is potent in focus reduction neutralization tests with authentic virus in Vero-TMPRSS2 cells. 2130-1-0114-112 potently neutralizes (A) WA1/2020 D614G (B) Delta B.1.617.2, (C) Omicron BA.1, (D) Omicron BA.1.1, (E) Omicron BA.2, (F) Omicron BA.2.12.1, (G) Omicron BA.4, (H) Omicron BA.5, and (I) Omicron BA.5.5 authentic viruses in focus reduction neutralization assays in Vero-TMPRSS2 cells. Symbols indicate the mean and standard deviation of two technical replicates; curves are 4-parameter logistic regression models fit of normalized data using GraphPad Prism. (J) IC50 values and 95% confidence intervals corresponding to (A)-(I). “>” indicates IC50 values > 10,000; “NC” indicates fits that were unconverged, unstable, or with positive hill slope. Analyses were performed in GraphPad Prism.

### Prophylaxis with 2130-1-0114-112 protects against SARS-CoV-2 variants

To assess the comparative prophylactic efficacy of 2130-1-0114-112 and the parental COV2-2130 mAb *in vivo*, we administered a single 100 μg (∼5 mg/kg total) dose to K18-hACE2 transgenic mice one day prior to intranasal inoculation with WA1/2020 D614G, BA.1.1, or BA.5 (88 mice in total, 9-10 for each mAb and viral strain). Although Omicron lineage viruses are less pathogenic in mice than humans, they replicate efficiently in the lungs of K18-hACE2 mice^27, 28^. Viral RNA levels were measured at 4 days post-infection in the nasal washes, nasal turbinates, and lungs (**Fig. 5**). As expected, the parental COV2-2130 mAb effectively reduced WA1/2020 D614G infection in the lungs (180,930-fold), nasal turbinates (42-fold) and nasal washes (25-fold) compared to the isotype control mAb. However, the COV2-2130 mAb lost protective activity against BA.1.1 in all respiratory tract tissues, whereas against BA.5, protection was maintained in the lungs (13,622-fold) but not in the nasal turbinates or nasal washes. Compared to the isotype control mAb (**Fig. 5**), 2130-1-0114-112 protected against lung infection by WA1/2020 D614G (399,945-fold reduction), BA.1.1 (53,468-fold reduction), and BA.5 (160,133-fold reduction). Moreover, in the upper respiratory tract (nasal turbinates and washes), 2130-1-0114-112 also conferred protection against WA1/2020 D614G, BA.1.1, and BA.5. The differences in protection between the parental COV2-2130 and derivative 2130-1-0114-112 mAbs were most apparent in mice infected with BA.1.1, which directly parallels the neutralization data (**Fig. 3 and 4**).

**Figure 5:**
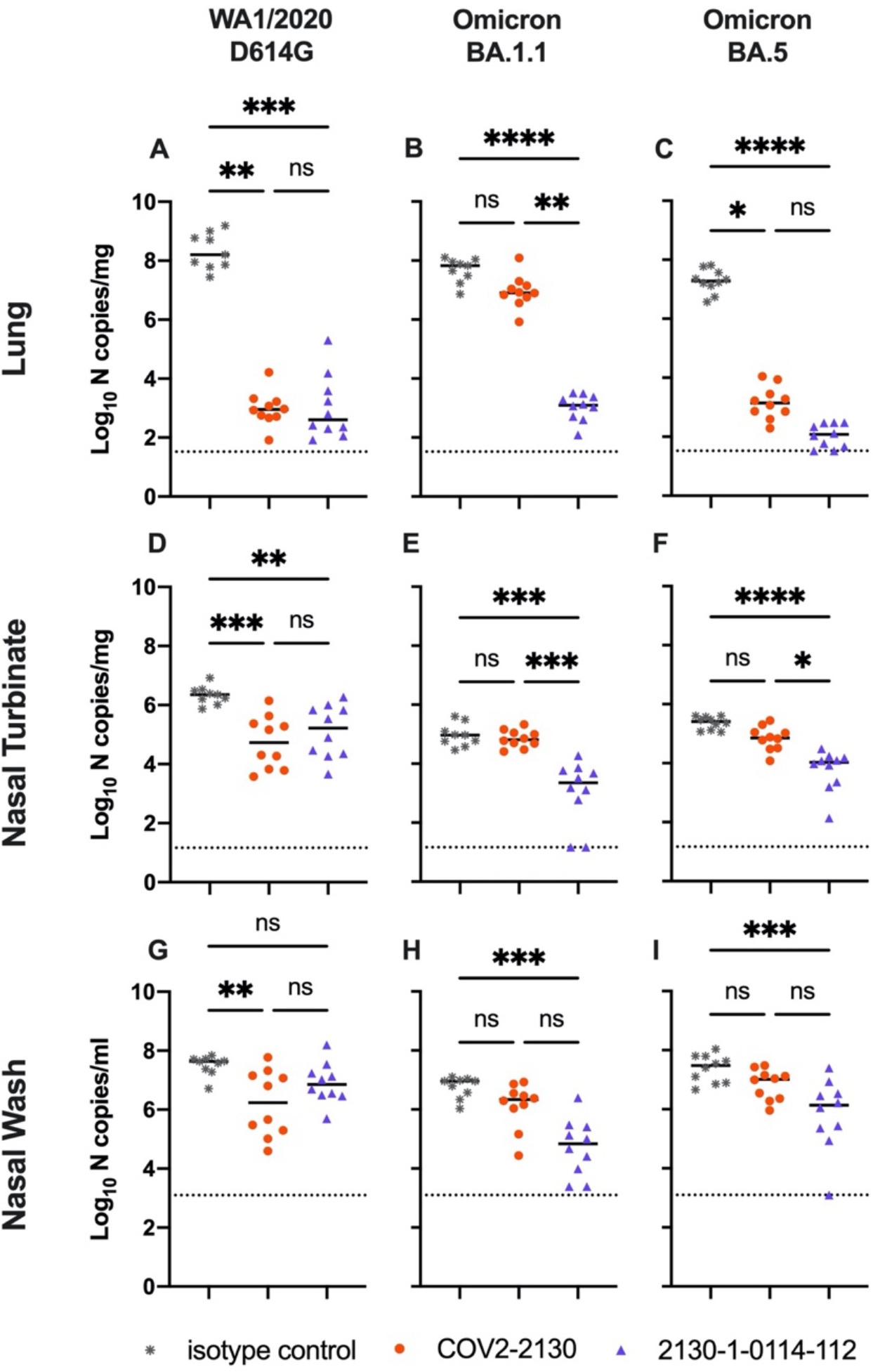
2130-1-0114-112 provides *in vivo* prophylactic protection against SARS-CoV-2 variants. Eight-week-old female K18-hACE2 mice were administered 100 μg (∼5 mg/kg) of the indicated mAb treatment by intraperitoneal injection one day before intranasal inoculation with 10^4^ FFU of WA1/2020 D614G (Left), Omicron BA.1.1 (Center) or BA.5 (Right). Tissues were collected four days after inoculation. Viral RNA levels in the lungs (Top), nasal turbinates (Center), and nasal washes (Bottom) were determined by RT-qPCR (lines indicate median of log_10_ values); n = 9 (WA1/2020 D614G and BA.1.1 isotype control groups) or 10 (all others) mice per group, two experiments; Kruskal-Wallis ANOVA with Dunn’s multiple comparisons post-test; ns, not significant; *P < 0.05, **P < 0.01, ***P < 0.001, ****P < 0.0001). All analyses conducted in GraphPad Prism.

### Deep mutational scanning reveals LLNL-112 improves broad potency without incurring additional escape liabilities

To understand the changes in 2130-1-0114-112 neutralization breath relative to its ancestral antibody, we next mapped the epitopes for both antibodies using spike-pseudotyped lentiviral deep mutational scanning^29^ (DMS). For each antibody we mapped escape mutations in both the BA.1 and BA.2 spikes. DMS experiments show that the escape profile of both COV2-2130 and 2130-1-0114-112 in the context of both BA.1 and BA.2 backgrounds is consistent with the epitope of the antibodies, but with differences in sensitivity to particular mutations (Fig. 6). Consistent with live and pseudovirus neutralization assays (Fig 3, Fig 4, **Fig ED3**), mutations at positions R346 and L452 are sites of significant escape from both antibodies (Fig. 6). In addition, both antibodies lose potency with mutations at site K444 (such as K444T found in BQ.1* variants). The reversion mutation S446G in the BA.1 background increases the neutralization potency of both antibodies (negative escape values in heatmaps) (Fig. 6C) and likely contributes to greater neutralization potency against the BA.2 variant (Fig. 3-4), which carries G446. Most mutations at sites K440 and R498 are slightly sensitizing to COV2-2130 antibody in both BA.1 and BA.2 backgrounds but provide weak escape for 2130-1-0114-112 in the BA.1 background and have even weaker or little effect in the BA.2 background. In agreement with pseudovirus neutralization assays (Fig. 3), comparison of mutation-level escape between the two antibodies shows that 2130-1-0114-112 antibody is substantially more potent than COV2-2130 against the BA.1 variant and retains better potency against viruses with additional mutations in both BA.1* and BA.2 backgrounds (Fig. 6 A-B). However, even with improved potency, 2130-1-0114-112 is still vulnerable to escape at multiple residues in the 444-452 loop, which is the site of convergent substitutions in several emerging Omicron lineages^30^. Many of these emerging variants contain multiple substitutions at positions identified by DMS as important for neutralization or in close proximity to the COV2-2130 epitope, including BQ.1.1 (R346T and K444T), XBB (R346T, V445P, G446S), and BN.1 (R346T, K356T, G446S). To understand the impact of these emerging VOCs, we assessed the ability of 2130-1-0114-112 to neutralize BQ.1.1, XBB, and BN.1 in pseudoviral neutralization studies. Consistent with the previously known liabilities of COV2-2130 and our deep mutational scanning results, 2130-1-0114-112 loses neutralizing activity against these VOCs (**Fig. ED4**), likely due to substitutions at 444 as well as combinatorial effects of multiple substitutions within the COV2-2130 epitope present in these variants. Taken together, these data demonstrate that 2130-1-0114-112 exhibits improved potency against many individual substitutions without incurring additional escape liabilities, although residues such as 444 remain critical for neutralization activity of both 2130-1-0144-112 as well as COV2-2130.

**Fig 6.**
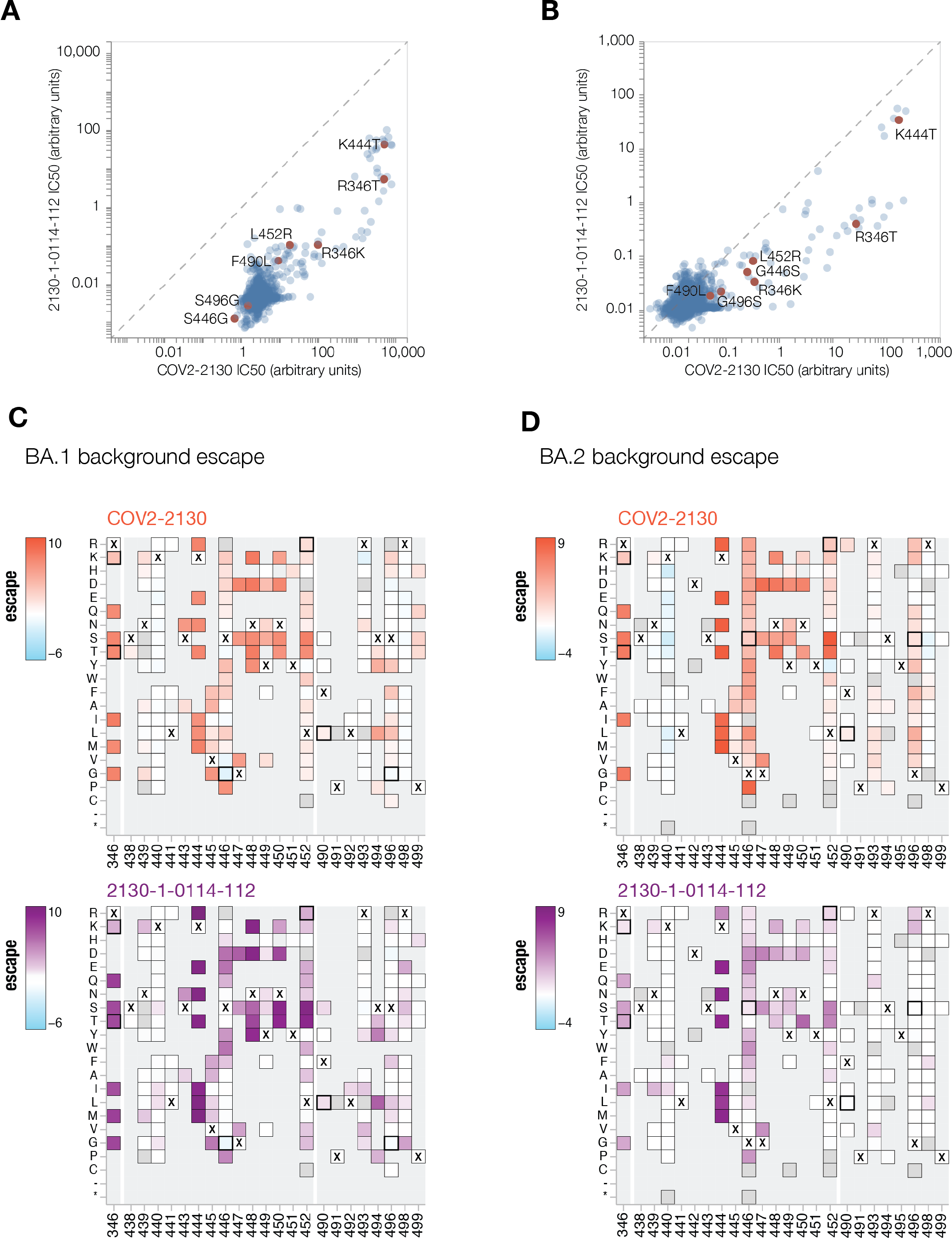
COV-2130 and 2130-1-0114-112 escape mapping using deep mutational scanning. **(A-B)** Comparison between IC50 values measured using deep mutational scanning for COV-2130 and 2130-1-0114-112 antibodies in (A) BA.1 and (B) BA.2 backgrounds with key mutations highlighted. Arbitrary units in both plots are on the same scale. Interactive plots that display each mutation can be found at https://dms-vep.github.io/SARS-CoV-2_Omicron_BA.1_spike_DMS_COV2-2130/compare_IC50s.html for BA.1 background and at https://dms-vep.github.io/SARS-CoV-2_Omicron_BA.2_spike_DMS_COV2-2130/compare_IC50s.html for BA.2 background. **(C-D)** Heatmaps of mutation escape scores at key sites for each antibody in (C) BA.1 and (D) BA.2 backgrounds. Escape scores are calculated relative to the wild-type amino acid in the same virus background. X marks wild-type amino acid in the relevant background. Amino acids not present in the deep mutational scanning libraries lack squares and gray squares are mutations that strongly impair spike-mediated infection. Mutations identified in (A) and (B) are shown with a heavy line surrounding the corresponding box. Interactive heatmaps for full spike can be found at for BA.1 background and https://dms-vep.github.io/SARS-CoV-2_Omicron_BA.2_spike_DMS_COV2-2130/COV2-2130_vs_2130-1-0114-112_escape.html for BA.2 background.

### Structural basis for the restored potency of 2130-1-0114-112

To elucidate the key intermolecular interactions that form the interface and determine Omicron RBD recognition by 2130-1-0114-112, we performed 3D reconstructions of the complex between SARS-CoV-2 Omicron BA.2 spike and 2130-1-0114-112 Fab fragment using cryo-electron microscopy (cryo-EM). Reconstruction using refinement of the full complex gave a map with average resolution of 3.26Å, but the interface region between the BA.2 RBD and 2130-1-0114-112 Fab was not well resolved, presumably due to the flexibility of the RBD-Fab region in the reconstruction. To resolve details at the intermolecular interface, we performed focused refinement of this portion of the structure. Focused refinement resulted in an effective resolution of ∼3.6Å for this region (EMD-28198, EMD-28199, PDB 8EKD) (**Fig. 7** and **Fig. ED7, Table EDT2**).

**Figure 7:**
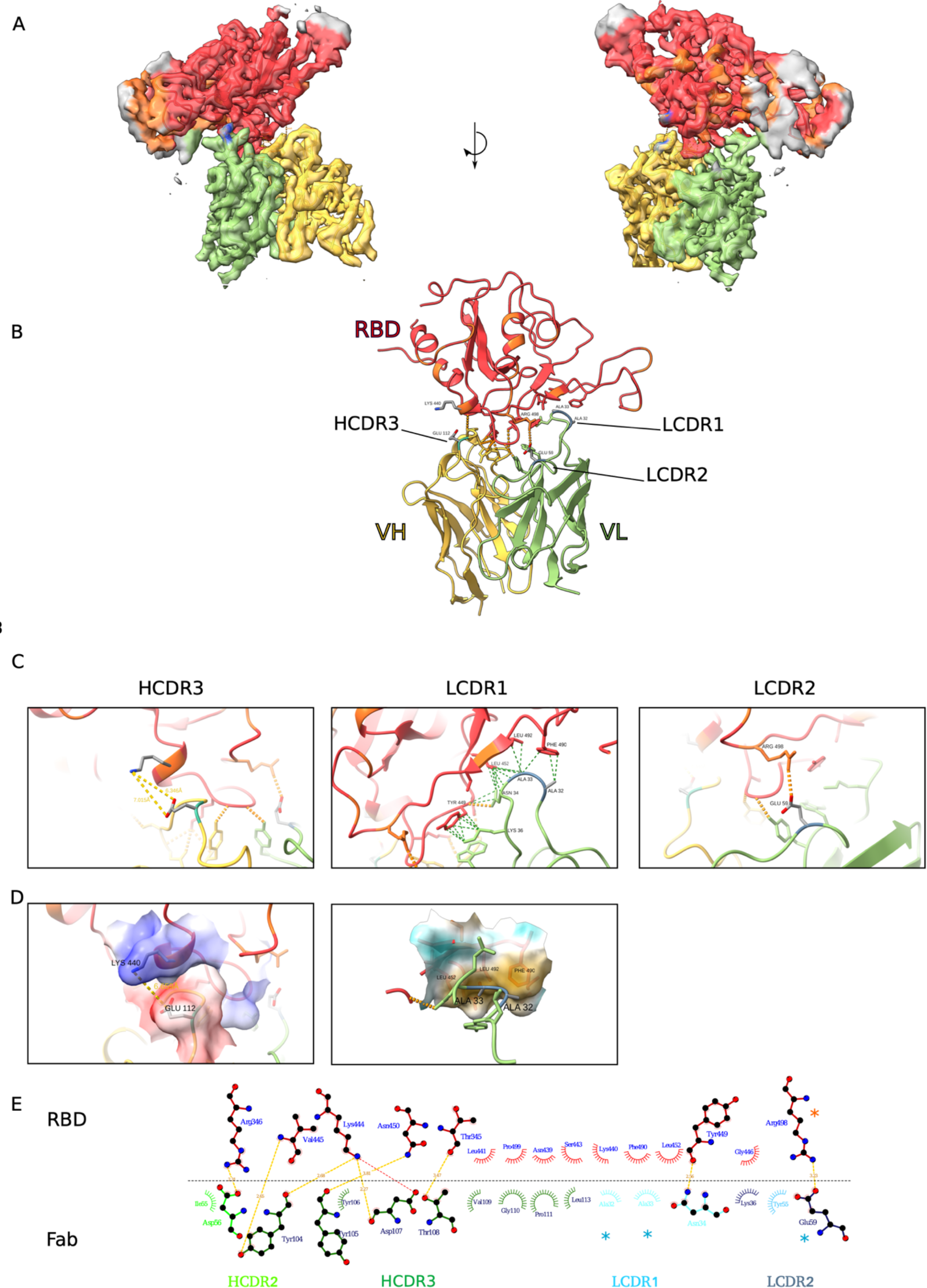
Cryo-EM structure of neutralizing antibodies 2130-1-0114-112 in complex with Omicron BA.2 RBD. (A) Cryo-EM map and model of the RBD-Fab complex. The map is transparent and colored by chain with RBD red, 2130-1-0114-112 HC yellow, and 2130-1-0114-112 LC green. (B) Atomic model of the RBD-Fab complex. Colors are the same as in A. Hydrogen bonds are shown as dashed lines. BA.2 RBD mutations are in orange. 2130-1-0114-112 mutation in cyan and blue (HC and LC). (C) Detail showing the 2130-1-0114-112 modified residues and the interaction with BA.2 RBD. Left, HCDR3 Glu112. Middle, LCDR1 Ala32 and Ala33 hydrophobic network. Right, LCDR2 Glu59 salt bridge with Arg498. Orange and green dashed lines indicate H-bonds and hydrophobic interactions, respectively; yellow dashed lines are labeled with distances. (D) Left, HCDR3 shown as in (C) with surface color by electrostatic potential, showing the positive and negative charges of Lys444 and Glu112. Right, A32 and A33 in LCDR1 with the nearby RBD surface colored by hydrophobicity (orange to cyan indicates hydrophobic to hydrophilic). (E) 2D diagram of Fab 2130-1-0114-112 paratope and epitope residues involved in hydrogen bonding (dashed lines) and hydrophobic interactions. Residues involved in hydrophobic interactions are shown as curved lines with rays. Atoms shown as circles, with oxygen red, carbon black, and nitrogen blue. Interacting residues that belong to CDR loops are colored in different shade. Asterisks correspond to mutated residues. Image created with Ligplot+^31^.

This model shows the binding interface of 2130-1-0114-112:RBD and elucidates how 2130-1-0114-112 regains neutralization potency against Omicron VOCs. The parental COV2-2130 forms extensive interactions with the RBD through HCDR2 and HCDR3, as well as LCDR1 and LCDR2^13^ with hydrogen bond networks and hydrophobic interactions. To improve binding interactions with Omicron subvariants, 2130-1-0114-112 modifies three CDR loops: G112E in HCDR3, S32A and S33A in LCDR1, and T59E in LCDR2.

The RBD N440K substitution, identified in the DMS as sensitizing for escape from COV2-2130 but less so for 2130-1-0114-112, is on the edge of the interface with the 2130-1-0114-112 HCDR3 loop and does not make direct contact with substitution G112E. However, N440K introduces a positive charge to a local environment that has substantial hydrophobic-to-hydrophobic contact. The negative charge introduced by the HCDR3 G112E substitution (**Fig. 7C, D**) might improve the electrostatic interactions in this region. It is possible that E112 and K440 are interacting by coordinating a water molecule but the structural resolution is not sufficient to confirm this type of interaction.

The local environment around the LCDR1 loop is mostly hydrophobic (comprised of RBD residues L452, F490 and L492, as well as the Omicron mutation E484A) with an N34 hydrogen bond (**Fig. 7D**). The hydrophilic-to-hydrophobic LCDR1 substitutions introduced in 2130-1-0114-112, S32A and S33A may favor the local environment and strengthen hydrophobic interactions with the RBD (**Fig. 7C, E**). This is supported by the DMS identification of sensitivity to hydrophobic-to-hydrophilic substitutions at RBD position 452 for both 1230-1-0114-112 and the parental COV2-2130. Lastly, the T59E mutation in the LCDR2 loop establishes a new salt bridge with the RBD substitution Q498R present in Omicron RBDs. This new salt bridge likely strengthens the interaction with the RBD (**Fig. 7C, E**).

2130-1-0114-112 distributes four substitutions across three of the four CDR loops comprising the parental COV2-2130 paratope. Mutations to CDRH3 loop were less fruitful than mutations in the L1 and L2 (**Fig. ED8** panel A in comparison to panel D) when looking across all antibody candidates. Among successful candidates, substitutions at positions 32 and 33 in CDRL1 appear enriched–particularly with hydrophobic residues–consistent with our analysis of this region of the experimentally solved structure of 2130-1-0114-112:BA.2 Spike. Another candidate, 2130-1-0104-024, achieves improved affinity and neutralization with only two substitutions, S32W in CDRL1 and T59E in CDRL2. However, full neutralization potency is not reached without the potential charge accommodation mediated by G112E. This suggests that a combination of new bonds and accommodating charge changes optimized the restored affinity and potency of 2130-1-0114-112 with Omicron variants. Altogether, the structural model of the 2130-1-0114-112 with the BA.2 RBD helps explain the observed restoration of potency against SARS-CoV-2 Omicron VOCs.

## Discussion

We set out to rapidly design and validate derivatives of the COV2-2130 antibody that restore potent *in vitro* neutralization against BA.1 and BA.1.1 Omicron subvariants while maintaining binding and neutralization to previous strains of SARS-CoV-2. Additionally, we sought to retain favorable thermostability properties and maintain the sequences’ humanness, a data-driven measure of similarity to known human sequences. Despite multiple mutations in the COV2-2130 epitope present in Omicron BA.1 and BA.1.1, we achieved these design objectives by applying a computationally driven, multi-objective approach. Several designed antibody candidates successfully restored neutralization potency to Omicron subvariants. In our top antibody design, 2130-1-0114-112, four substitutions accommodate Omicron escape mutations in BA.1 and BA.1.1 without sacrificing potency against Delta. This engineered antibody is thermostable and also potently neutralizes Omicron BA.2, BA.4, BA.5, and BA.5.5. Further, this antibody also displayed restored prophylactic efficacy *in vivo*. This work demonstrates our approach for extending the utility of a high-value antibody, complementing state-of-the-art *ex vivo* antibody discovery of such high-value antibodies with responsive computational modification.

It is plausible that the distributed nature of the improvements in 2130-1-0114-112, with four mutations spanning three CDRs, makes this antibody comparatively robust to subsequent escape. This supposition is supported by deep mutational scanning results that demonstrate an improvement in binding over the parental COV2-2130 where escape substitutions present in the design targets, BA.1 and BA.1.1, are mitigated without incurring any new sites of vulnerability. 2130-1-0114-112 does not mitigate the parental COV2-2130 antibody’s reliance on K444 and sensitivity to substitutions at this residue. Although our top candidate does not neutralize the most recent variants BQ.1.1 and XBB, which contain multiple substitutions within the COV2-2130 epitope, deep mutational scanning results indicate that 2130-1-0114-112 reduces the impact of some of the individual mutations present in these variants.

Our design approach could potentially expedite the path of new drug products to clinical use, including lower development costs and lower risk as compared to identifying a wholly new drug product of comparable breadth and efficacy. Our top performing antibody restores *in vivo* efficacy and achieves potent and broad neutralization of many SARS-CoV-2 VOCs by substituting only four amino acids into the parental antibody. This parental antibody has previously been extensively tested for safety, manufacturability, and clinical efficacy^1^. Given increasing evidence that neutralization is a correlate of protection from severe COVID-19 in patients treated with monoclonal antibody therapies^32, 33^, an immunobridging strategy has been proposed as a response to rapidly evolving SARS-CoV-2 variants to shorten the pathway of new monoclonal antibodies to clinical use, particularly those based on existing products with demonstrated efficacy and safety in clinical trials^34^. Rapid computational rescue of high-value, potentially rare, antibodies in clinical use is a direct application of our work and, combined with a future immunobridging strategy, is particularly relevant now given that existing antibody drug development approaches are currently struggling to match the rapid pace of SARS-CoV-2 evolution.

While the individual components comprising our approach are built upon existing computational approaches, we integrate them into a novel framework that demonstrates (1) a computational approach to antibody optimization that gains neutralization to a new target (i.e., the BA.1.1 escape variant for which there was complete or near-complete loss of COV2-2130 neutralization activity), (2) successful optimization of an antibody to achieve high potency to multiple targets (e.g., multiple escape variants) without requiring experimental iterations, and (3) computationally restoring or improving efficacy with *in vivo* validation. The computational approach we used in this work did not require iterative improvement based on feedback from experimental evaluations, nor did it require availability of data on antibody candidates tested against the target antigens, either of which would result in further delays when responding to a newly emerged variant.

In future work, we will extend our computational approach to include additional antibody developability predictive models, such as models predicting antibody expression, protein aggregation, and polyreactivity. Our models for predicting antibody-antigen binding heavily depend on performing simulations with sufficiently accurate models of antibody-antigen co-structures, an important limitation. Consequently, we are developing experimental datasets to advance machine learning-based approaches for predicting binding directly from sequence, as well as incorporating emerging artificial intelligence-based approaches for determining and refining structural models.

In summary, we demonstrate critical aspects of an antibody design capability and rapidly create hundreds of antibody designs, some of which are potently neutralizing and broadly reactive replacement antibodies for COV2-2130 in the context of Omicron subvariants. Such an approach could lead to an on-demand antibody drug product development strategy that would allow for rapid response to emerging viral outbreaks as well as in response to viral evolution. Leveraging additional technologies, such as deep mutational scanning approaches we employ in this work, together with a computational capability to co-optimize antibodies to multiple targets and mutational liabilities, could transform the process of antibody discovery and enable the pre-emptive optimization of antibodies with increased robustness to escape.

## Methods

### Problem formulation: Generating antibody-antigen co-structures

To best manage the high sensitivity of protein binding affinity (herein considered as mutational changes to the binding free energy, ddG) predictions to antibody–antigen structure quality^35^, we used the program LGA^36^ to evaluate compatibility between numerous experimentally solved structures of the receptor binding domains (RBD), available structures of the Fab form of COV2-2130, and structures of RBD-Fab complexes. This approach allowed us to identify regions of backbone and side-chain deviation (see **Fig. ED9A**).

We used the conformational centroid to select a representative complex for further analysis. Structural clustering of tested RBDs identified Omicron RBD (PDB id 7t9k, chain A) as the centroid of all evaluated conformations (shown on **Fig. ED9B**). We consequently chose to perform ddG calculations on two initial structures (**Fig. ED9C**): an experimentally solved structure of WT RBD with the Fab form of COV2-2130 (PDB ID 7l7e, chains S, M, N), and a structural model of Omicron RBD complexed with COV2-2130 (PDB ID 7l7e, chains M, N) that uses the RBD as the identified conformational centroid (PDB ID 7t9k chain A).

### Problem formulation: Defining the search space

We specify which antibody positions to consider for mutation based on our estimated co-structures. We consider a given position for mutation if its wild-type residue includes any atoms less than 7 Å from any antigen atom. Under this criterion, we consider 25 positions for mutation. For each position considered, we allow amino acid substitutions to all amino acids except cysteine or proline. We limit each mutant sequence to a maximum of 9 amino acid substitutions relative to wild-type COV2-2130. This results in a search space of size 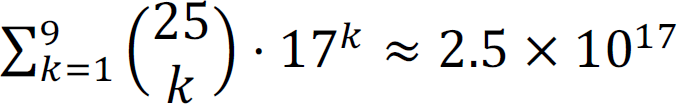.

### Semi-autonomous system

In the two sections below, we describe how our semi-autonomous system proposes mutant antibody sequences and selects which sequences to simulate via Rosetta Flex^18^. Decision-making agents run in parallel in an asynchronous, distributed fashion. Each agent operates on a single HPC node and selects a number of sequences to simulate according to the number of available cores. Simulation results (e.g., Rosetta Flex ddG calculations) are recorded in a centralized database so all agents have access to all results. Initially, the system simulates all single-point mutant sequences using MD, SFE, FEP, Rosetta Flex, and FoldX, as well as scores them under AbBERT.

### Sequence generation

Mutant antibody sequences are proposed using a sequence generator that operates using a hierarchical sampling process. First, the number of mutations is uniform randomly sampled from 1 to 8. Then, mutations are sampled without replacement according to a probability distribution informed by six tools: Atomistic MD, SFE, FEP, Rosetta Flex, FoldX, and AbBERT. Concretely, each tool outputs a score (e.g., -ddG for MD) for all single-point mutant sequences. Each score is then converted into an unnormalized mutation probability by passing it through a generalized logistic function:

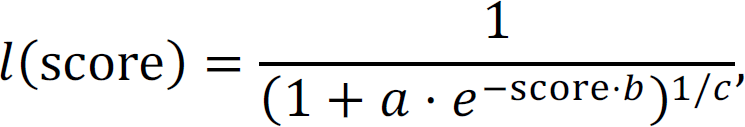

where *a* = 1000, *b* = 5, and *c* = 2 were hand-tuned to strongly prefer positive scores and to effectively squash or truncate extreme values (e.g., ddG’s of -3 and -3.5 produce very similar scores). Finally, we obtain a normalized probability distribution by normalizing across all tools and all possible mutations. Thus, the probability of mutation *m* is given by:

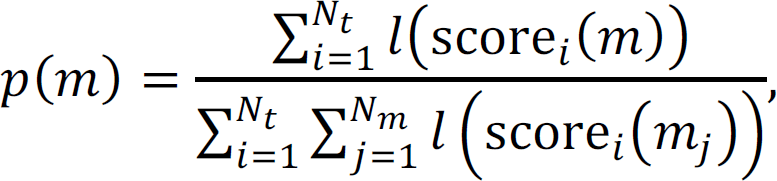

where *N_t_* is the number of tools, *N_m_* is the number of possible mutations, score_*i*_ is the score of the *i*th tool, and *m_j_* is the *j*th possible mutation.

### Sequence selection

We employ Bayesian optimization agents to select which sequences to simulate via Rosetta Flex. Within the optimization loop, we first generate batches of sequences from our sequence generator. We then use Gaussian process (GP) agents, described below, to estimate the posterior distribution for each proposed sequence and select a subset of sequences according to the maximum expected improvement (MEI) acquisition function^37^. Finally, the subset of sequences are simulated using Rosetta Flex, and the optimization loop continues.

The input to the GP is based on a count-based featurization of the mutant antibody sequence and a multilayer perceptron (MLP). The count-based featurization is based on chemical and size properties of mutant sequences, described as follows. In the starting, unmutated co-structure, we identify pairs of antibody-antigen amino acids with α-carbon to α-carbon distances less than 10 Å. This establishes a bipartite graph of antibody and antigen amino acids, where the vertices are antibody and antigen amino acids and the edges are the identified antibody-antigen amino acid pairs. Given a mutant sequence, we substitute amino acid mutations at the vertices without altering the graph structure. For each vertex (amino acid), we assign one or more chemical properties (acidic, aliphatic, aromatic, basic, hydroxylic, sulfuric); see **Table 1**. For each edge (antibody amino acid-antigen amino acid), according to the two connected vertices, we assign one or more unordered pairs of chemical properties (e.g., acidic-sulfuric, basic-basic), of which there are 28 possible pairs. Similarly, we assign each vertex a size property (very small, small, medium, large, very large; see **Table 1**) and each edge an unordered pair of size properties (e.g., small-medium), of which there are 15 possible pairs. We define the first 43 features as the counts of each chemical and size property pair. The second 43 features are these same counts when using wild-type antibody.

**Table 1.** Amino acid chemical and size classifications used in the count-based featurization; modified from Pommié, 2004^38^.

| *Size\Chemical Class* | Aliphatic | Aromatic | Acidic | Basic | Hydroxilic | Sulfuric | Amidic |
| --- | --- | --- | --- | --- | --- | --- | --- |
| *Very Large* |  | F, W, Y |  |  |  |  |  |
| *Large* | I, L |  |  | K, R |  | M |  |
| *Medium* | V |  | E | H |  |  | Q |
| *Small* | P |  | D |  | T | C | N |
| *Very Small* | A, G |  |  |  | S |  |  |

The resulting 86-dimensional feature vector is then used as input to a multilayer perceptron (MLP), comprising a single hidden layer with output dimension 40 and tanh activation, followed by an output layer with output dimension 10 and no activation. The output of the MLP is then used as the input to the GP.

The GP is defined by a mean and kernel covariance function *GP*(μ(*x*), μ(*x*, *x*’)), a non-zero constant prior mean function μ(*x*) = *C*, where *C* is a learned scalar parameter, and a scaled radial basis function kernel operating on inputs *x*_1_ and *x*_2_:

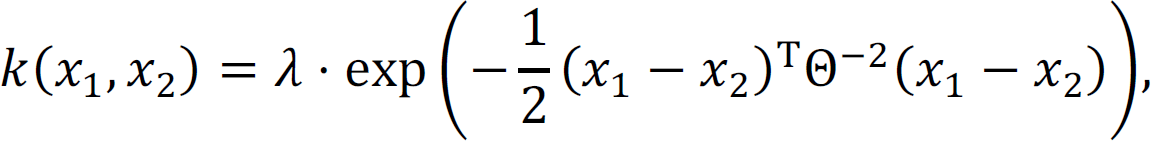

where Θ is a length scale parameter and λ is a kernel scale parameter. GP parameters (*C*, Θ, λ) and MLP parameters were jointly trained using stochastic gradient descent on the log marginal likelihood of Rosetta Flex ddG values of 20,000 training sequences from the sequence generator with respect to the GP likelihood function, *p*(*y* ∣ *X*).

We compute the GP predictive posterior distribution *f*_∗_ as *p*(*f*_∗_ ∣ *X*_∗_, *X*, *y*), where *X* is the union of the 20,000 training sequences and 10,000 sequences randomly drawn from the centralized database (all passed through the featurization and MLP), and *y* represents the union of the corresponding Rosetta Flex ddG values. To select a batch of sequences, we first generate 1,000 candidate sequences, *X*_∗_, from the sequence generator. We compute marginal posteriors for each sequence, then select the sequence with the maximum expected improvement, where we are minimizing the free energy:

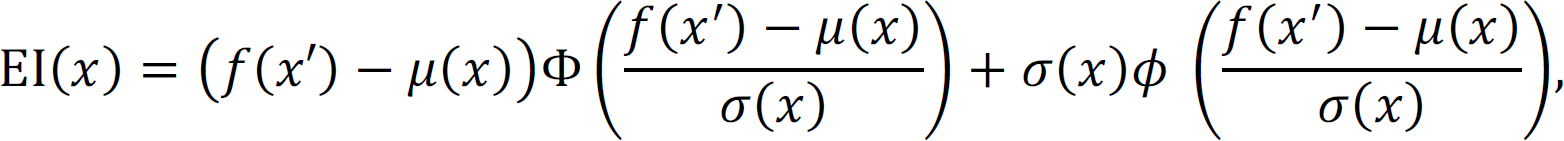

where *x* is the candidate sequence, *x*’ is the sequence with lowest ddG acquired so far, μ(x) is the mean of the GP predictive posterior at *x*, σ(*x*) is the standard deviation of the GP posterior at *x*, and Φ and ϕ are the cumulative density and probability density functions of the normal distribution. Upon selection, *X* and *y* are updated using the selected sequence and the predicted ddG, respectively. Then the predictive posterior *f*_∗_ is re-computed. Subsequently, we select the next candidate according to MEI; this process continues until enough sequences are selected to occupy all cores.

To supplement the sequences chosen by the Bayesian optimization agents, and to ensure sufficient coverage in sequence space, we employ two additional types of rules-based agents. The first agent simply selects all combinations of two-point mutant sequences. The second agent samples from among the current top-performing sequences (i.e., those with the most negative ddG) and further mutates them according to the sequence generator described above.

### Down-selecting sequences for experimental validation

The autonomous system described above produced over 125,000 sequences simulated using Rosetta Flex. To down-select this set to our experimental capacity of 376 candidate sequences, we first computed the Pareto (non-dominated) set of sequences, based on the objectives listed in **Table 2**. Note that MD, SFE, and FEP multi-point mutation scores were approximated as the sum of their constituent single-point mutation scores; multi-point AbBERT scores were computed on all sequences. The resulting Pareto set contained 3,809 sequences. From here, we sought consensus across tools by ranking all sequences in the Pareto set according to the weighted sum of each objective, with a penalty based on mutational distance from wild-type COV2-2130:

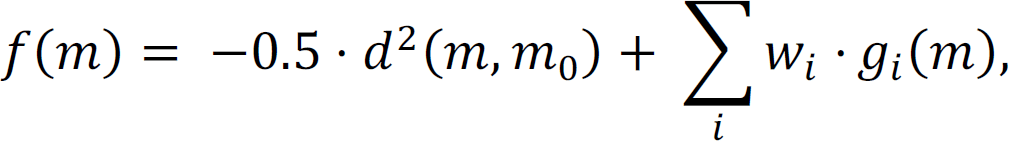

**Table 2.** Objectives considered for Pareto selection of candidate sequences, with associated units and weights.

| Objective ($g_i$) | Units | Weight ($w_i$) |
| --- | --- | --- |
| Rosetta Flex ddG, L452R (Delta) | Rosetta Energy Units (REU) | -1.0 |
| Rosetta Flex ddG, BA.1 | REU | -1.0 |
| Rosetta Flex ddG, BA.1.1 | REU | -1.0 |
| Atomistic MD ddG BA.1 | KCal/Mol | -1.5 |
| SFE ddG, BA.1 | REU | -1.5 |
| SFE ddG, BA.1.1 | REU | -1.5 |
| FoldX ddG, L452R (Delta) | KCal/Mol | -0.5 |
| FoldX ddG, BA.1 | KCal/Mol | -0.5 |
| FoldX ddG, BA.1.1 | KCal/Mol | -0.5 |
| FEP stability ddG | KCal/Mol | -0.5 |
| AbBERT score | Arbitrary Units (AU) | 2.0 |

where *m* is a mutant sequence, *m*_9_ is wild-type COV2-2130, *g_i_* is the *i*th objective, *w*_i_ is the *i*th weight, and *d* is the count of amino acid substitutions relative to wild-type COV2-2130.

We expect that some simulations will systematically misestimate the value of some mutations. Enforcing sequence diversity of selected antibodies may mitigate risk of systematic errors in our simulation tools. Thus, we enforced sequence diversity when selecting among the top-ranked sequences. First, we limited the number of times any particular mutation could appear in the final set; lower-ranked sequences beyond this limit were excluded from selection. Second, for each tool, we enforced inclusion of at least one sequence containing that tool’s top-performing single-point mutations, even if disfavored by other tools. Third, to ensure mutational diversity across positions, we enforced inclusion of at least one sequence containing a mutation at interface positions, even if not scored favorably by the tools. Finally, we excluded sequences containing more than four mutations to aromatic residues and sequences containing glycosylation motifs. After eliminating enforced exclusions from the ranked list and selecting enforced inclusions, we selected remaining top-ranked sequences.

### Atomistic molecular dynamics (MD) simulations for free energies as affinity predictions

We performed MD simulations using OpenMM (v7.4)^39^ and CHARMM36^40^. Complexes were first solvated in an isotropic TIP3P^41^ box. K^+^ and Cl^-^ ions were then added to neutrality and 150 mM concentration. After energy minimization, we ran MD simulations with a Langevin integrator (1 ps^-1^)^42^. Monte Carlo barostat (303.15 K), particle mesh ewald summation (1 Å grid)^43^, and SHAKE^44^. Simulations proceeded in 2 fs timesteps for a total of 125 ps with constraints on backbone and sidechain atoms (400 and 40 kJ/mol·nm^2^, respectively). An additional 10 ns were then run without constraints.

From the final coordinates, we increased sampling using a minimum watershell^45^ with adaptive boundary and hydrogen masses were increased to 4 amu by transferring the mass from the bonded non-hydrogen atom. These simulations employed a 4 fs time step^46^, 300K thermostat, and particle mesh ewald electrostatics. The antibody and antigen were separated by 8 Å, with individual simulations under harmonic constraints (100 kcal/mol·Å^2^) at 1 Å intervals. At each 1 Å interval, we ran 4 ns of re-equilibration and an additional 320 ns of MD to provide sampling needed to calculate the free energy^47^. Sampling of the CDR loops might benefit from using a force field tuned to reproduce conformations of intrinsically unstructured proteins^48^.

### Structural Fluctuation Estimation (SFE) approach for reproducible and robust free energy prediction

We applied our Structural Fluctuation Estimation (SFE) approach^17^ to address problems of reproducibility and robustness of calculated estimates in energy changes upon mutations (ddG). Antibody-antigen structures were minimized and relaxed using standard minimization procedures from Rosetta^49^, Chimera^50^, and GROMACS^51^ steepest descent and conjugate gradient methods; short MD simulations in GROMACS subsequently extracted a set of structure snapshots from the resulting trajectories.

For each initial structure, we generated 60 structural conformations for the RBD-Fab complex— 30 complexes with mutations, and 30 without. Each set of 30 complexes includes the initial structure, 4 minimized structures, and 25 structures from MD trajectories, so as to capture structural uncertainties, possible structural deviations upon introduced mutations, and natural fluctuations in protein structure. Using Rosetta Flex ddG^18^, we performed mutational ddG calculations in the “forward” direction on models without mutations, and in “reverse” on models with mutations. Once ddG calculations were completed, we removed outliers, averaged results of the interquartile simulations, and calculated the final ddG estimate via the formula: ddG = (ddG_forward_ - ddG_reverse_) / 2. The resulting ddG value provides an affinity estimate shown to be more reproducible and robust than ddG estimates calculated from only one initial input structure of the RBD-Fab complex, whether using standard FoldX^19^, Rosetta^52^, or Flex ddG^18^ procedures.

### Free energy perturbation calculations

Free energy perturbation (FEP) is a rigorous, physics-based method for calculating free energy differences that employs MD simulations. As reported recently^20^, we implemented an automated protocol for large-scale FEP calculations to evaluate the effect of antibody mutation on conformational stability. The structure of the COV2-2130 Fab was taken from the crystal structure 7L7E. Using the FEP protocol described^20^, we calculated the change in antibody conformational stability for all single-point mutations. We first considered an extended set of 29 residues to assess whether COV2-2130 exhibited stability liabilities outside the 25 residues described above. We subsequently limited mutations to only the 25 positions in the multi-point optimization.

### AbBERT language model

AbBERT^21^ is a transformer-based language model derived by fine-tuning the pre-trained ProtBERT^53^ language model on over 200,000 human antibody sequences obtained from the Observed Antibody Space (OAS) database^22^. The trained AbBERT model estimates the distribution of human antibody sequences, providing a way to measure the resemblance of candidate antibodies to human antibodies. We scored the humanness of mutant sequences via a multi-unmask scoring procedure. **Fig. ED10** illustrates AbBERT unmasking probabilities for the CDRs of COV2-2130.

### Antigen production

To express the RBD subdomain of the SARS-CoV-2 S protein, residues 328–531 were cloned into a mammalian expression vector downstream of a mu-phosphatase signal peptide and upstream of an AviTag and a 8×His tag. Three previously identified stabilizing mutations (Y365F, F392W, V395I) were included in the RBD to enhance antigen stability and yield. For RBD constructs corresponding to the Omicron subvariants, mutations present in each subvariant were introduced into the context of the stabilized, wild-type RBD construct. RBD constructs were transfected into Expi293F cells (ThermoFisher Scientific), and expressed protein was isolated by metal affinity chromatography on HisTrap Excel columns (Cytiva). For structural studies, we used a previously described stabilized SARS-CoV-2 spike construct (VFLIP). This construct contains an inter-protomer disulfide bond, a shortened linker between the S1 and S2 domains, and five proline substitutions relative to the native sequence of SARS-CoV-2 spike. In addition to these modifications, this construct also contains a c-terminal T4 fibritin foldon domain as well as an 8×His tag and a TwinStrep tag for purification. To express the protein, we transfected Expi293F cells (ThermoFisher Scientific) with a plasmid encoding SARS-CoV-2 S_VFLIP with BA.2 amino acid substitutions. Culture supernatants were collected 4-5 d after transfection and clarified by centrifugation. BioLock (IBA Biosciences) was added to remove free biotin in the culture media, after which supernatants were filter-sterilized using a 0.2 µm filter. Full-length VFLIP_BA.2 was purified by streptactin affinity chromatography using StrepTrap HP columns (Cytiva) and eluted using an elution buffer of 25 mM desthiobiotin in Dulbecco’s phosphate-buffered saline (DPBS). After elution, spike protein was prepared for use in cryo-electron microscopy (Cryo-EM) by size-exclusion chromatography. Purified proteins were analyzed by SDS-PAGE to assess purity and appropriate molecular weights.

### Antibody production

For each antibody in the first set of 230 designs, nucleotide sequences encoding the designed heavy and light chain sequences were synthesized, cloned into an hIgG1 framework, and used to produce mAbs via transient transfection of HEK293 cells at ATUM (Newark, CA, USA).

For the second set of 204 designs, monoclonal antibody sequences were synthesized (Twist Bioscience), cloned into an IgG1 monocistronic expression vector^54^ (designated as pVVC-mCisK_hG1), and expressed either at microscale in transiently transfected ExpiCHO cells^55^ for screening, or at larger scale for down-stream assays. Sequences in this group of 204 designs all contain an additional arginine at the beginning of the light chain constant region with respect to sequences expressed in the first set. Larger-scale monoclonal antibody expression was performed by transfecting (30 ml per antibody) CHO cell cultures using the Gibco ExpiCHO Expression System and protocol for 125ml flasks (Corning) as described by the vendor. Culture supernatants were purified using HiTrap MabSelect SuRe (Cytiva, formerly GE Healthcare Life Sciences) on a 24-column parallel protein chromatography system (Protein BioSolutions). Purified monoclonal antibodies were buffer-exchanged into PBS and stored at 4 °C until use.

### Binding screening and characterization

Immunoassays for screening the first set of 230 designs (**Fig. ED1**) and later characterization were performed on the Gyrolab xPlore instrument (Gyros Protein Technologies) using the Bioaffy 200 discs (Gyros Protein Technologies). The standard manufacturer’s immunoassay automated protocol was executed with fluorescence detection set to 0.1% PMT. Assay column washes were performed in PBS + 0.02% Tween 20 (PBST). Capture antigens were applied to the assay columns at 0.5 to 2.0 μM in PBS. Analyte mAbs were applied to the assay columns diluted in PBST at 1:200 for single-concentration screening or as a serial dilution from 1,000 nM to 0.25 nM for characterization of down-selected candidate antibodies. A secondary detection antibody served as a fluorescent reporter: Alexa Fluor 647 AffiniPure Fab Fragment Goat Anti-Human IgG, Fcγ fragment specific (Jackson ImmunoResearch) diluted to 50-100 nM in RexxipF buffer (Gyros Protein Technologies). Resulting values were fit to a 4PL model or calculated as area under the curve (AUC) using GraphPad Prism software.

### Dose-response ELISA binding assays

For screening and characterizing the second set of 204 designs (**Fig. ED2**), wells of 384-well microtiter plates were coated with purified recombinant SARS-CoV-2 RBD proteins at 4 °C overnight at an antigen concentration of 2 mg/mL. Plates were washed with Dulbecco’s phosphate-buffered saline (DPBS) containing 0.05% Tween-20 (DPBS-T) and blocked with 2% bovine serum albumin and 2% normal goat serum in DPBS-T (blocking buffer) for 1 h. mAbs were diluted in 12 three-fold serial dilutions in blocking buffer at a starting concentration of 10 µg/mL. Plates were then washed and mAb dilutions were added and incubated for 1 h. Plates were washed, a goat anti-human IgG conjugated with horseradish peroxidase (HRP) (Southern Biotech, cat. 2014-05, lot L2118-VG00B, 1:5,000 dilution in blocking buffer) was added, and the plates were incubated for 1 h. After plates were washed, signal was developed with a 3,3’,5,5’-tetramethylbenzidine (TMB) substrate (Thermo Fisher Scientific). Color development was monitored, 1M hydrochloric acid was added to stop the reaction, and the absorbance was measured at 450 nm using a spectrophotometer (Biotek). Dose-response ELISAs were performed in technical triplicate with at least two independent experimental replicates.

### Thermal Shift Protein Assays (melt-curve assays)

Antibody concentrations were determined using the Qubit Protein Assay Kit (ThermoFisher). The GloMelt™ Thermal Shift Protein Stability Kit (Biotum) was utilized to determine the thermal stability of the antibodies by following the manufacturer’s suggested protocols. The analysis was performed using a melt-curve program on an ABI 7500 Fast Dx Real-Time PCR instrument. Each assay was done in triplicate, using 5ug of mAb per well. The raw melt curve data was imported into and analyzed via Protein Thermal Shift ™ software version 1.4 (ThermoFisher) to generate the melting temperature and fit data.

### Pseudovirus Neutralization

Pseudovirus neutralization assays were carried out according to the protocol of Crawford *et al*^56^. One day prior to the assay, 293T cells stably expressing human ACE2 (293T-hACE2 cells) were seeded onto 96-well tissue culture plates coated with poly-D-lysine. The day of the assay, serial dilutions of monoclonal antibodies in duplicate were prepared in a 96-well microtiter plate and pre-incubated with pseudovirus for 1 h at 37 °C in the presence of a final concentration of 5 mg/mL polybrene (EMD Millipore), before the pseudovirus-mAb mixtures were added to 293T-hACE2 monolayers. Plates were returned to the 37 °C incubator, and then 48-60 h later luciferase activity was measured on a CLARIOStar plate reader (BMG LabTech) using the Bright-Glo Luciferase Assay System (Promega). Percent inhibition of pseudovirus infection was calculated relative to pseudovirus-only control. IC50 values were determined by nonlinear regression using Prism v.8.1.0 (GraphPad). Each neutralization assay was repeated at least twice.

### Viruses: FRNT and in vivo protection

The WA1/2020 recombinant strain with D614G substitution and B.1.617.2 was described previously^28, 57^. The BA.1 isolate was obtained from an individual in Wisconsin as a mid-turbinate nasal swab^58^. The BA.1.1 and BA.2 strains were obtained from nasopharyngeal isolates. The BA.2.12.1, BA.4, BA.5, and BA.5.5 isolates were generous gifts from M. Suthar (Emory University), A. Pekosz (Johns Hopkins University), and R. Webby (St. Jude Children’s Research Hospital). All viruses were passaged once on Vero-TMPRSS2 cells and subjected to next-generation sequencing^59^ to confirm the introduction and stability of substitutions. All virus experiments were performed in an approved biosafety level 3 (BSL-3) facility.

### Focus Reduction Neutralization Test

Serial dilutions of sera were incubated with 10^2^ focus-forming units (FFU) of WA1/2020 D614G, B.1.617.2, BA.1, BA.1.1, BA.2, BA.2.12.1, BA.4, BA.5, or BA.5.5 for 1 h at 37°C. Antibody-virus complexes were added to Vero-TMPRSS2 cell monolayers in 96-well plates and incubated at 37°C for 1 h. Subsequently, cells were overlaid with 1% (w/v) methylcellulose in MEM. Plates were harvested 30 h (WA1/2020 D614G and B.1.617.2) or 70 h (BA.1, BA.1.1, BA.2, BA.2.12.1, BA.4, BA.5, and BA.5.5) later by removing overlays and fixed with 4% PFA in PBS for 20 min at room temperature. Plates were washed and sequentially incubated with a pool (SARS2-02, -08, -09, -10, -11, -13, -14, -17, -20, -26, -27, -28, -31, -38, -41, -42, -44, -49, -57, -62, -64, -65, -67, and -71)^60^ of anti-S murine antibodies (including cross-reactive mAbs to SARS-CoV) and HRP-conjugated goat anti-mouse IgG (Sigma Cat # A8924, RRID: AB_258426) in PBS supplemented with 0.1% saponin and 0.1% bovine serum albumin. SARS-CoV-2-infected cell foci were visualized using TrueBlue peroxidase substrate (KPL) and quantitated on an ImmunoSpot microanalyzer (Cellular Technologies).

### Mouse studies

Animal studies were carried out in accordance with the recommendations in the Guide for the Care and Use of Laboratory Animals of the National Institutes of Health. The protocols were approved by the Institutional Animal Care and Use Committee at the Washington University School of Medicine (assurance number A3381–01). Virus inoculations were performed under anesthesia that was induced and maintained with ketamine hydrochloride and xylazine, and all efforts were made to minimize animal suffering. Heterozygous K18-hACE2 C57BL/6J mice (strain: 2B6.Cg-Tg(K18-ACE2)2Prlmn/J, Cat # 34860) were obtained from The Jackson Laboratory. Animals were housed in groups and fed standard chow diets.

Eight-week-old female K18-hACE2 C57BL/6 mice were administered 100 μg of 2130-1-0114-112, parental 2130, or isotype control anti-West Nile virus hE16 mAb^61^ by intraperitoneal injection one day before intranasal inoculation with 104 focus-forming units (FFU) of WA1/2020 D614G, BA.1.1 or BA.5. Animals were euthanized at 4 days post-infection and tissues were harvested for virological analysis.

### Measurement of Viral RNA burden

Tissues were weighed and homogenized with zirconia beads in a MagNA Lyser instrument (Roche Life Science) in 1 ml of DMEM medium supplemented with 2% heat-inactivated FBS. Tissue homogenates were clarified by centrifugation at 10,000 rpm for 5 min and stored at −80°C. RNA was extracted using the MagMax mirVana Total RNA isolation kit (Thermo Fisher Scientific) on the Kingfisher Flex extraction robot (Thermo Fisher Scientific). RNA was reverse transcribed and amplified using the TaqMan RNA-to-CT 1-Step Kit (Thermo Fisher Scientific). Reverse transcription was carried out at 48°C for 15 min, followed by 2 min at 95°C. Amplification was accomplished over 50 cycles as follows: 95°C for 15 s and 60°C for 1 min. Copies of SARS-CoV-2 *N* gene RNA in samples were determined using a published assay^62^.

### Plaque Assay Neutralization Tests

All SARS-CoV 2 viral stocks and VAT cells used for plaque assays were obtained through BEI Resources, NIAID, NIH. Delta variant (isolate hCoV-19/USA/MD-HP05647/2021, lineage B.1.617.2; NR-55672) was contributed by Dr. Andrew S. Pekosz. BA.1 (isolate hCoV-19/USA/GA-EHC-2811C/2021, lineage B.1.1.529; NR-56481) was contributed by Mehul Suthar. BA1.1 (isolate hCoV-19/USA/HI-CDC-4359259-001/2021, lineage B.1.1.529; NR-56475) was contributed by Centers for Disease Control. Viral stocks were amplified in Vero E6 cells (Delta variant) or VAT cells (Omicron variants). Serial dilutions of mAbs were incubated with virus at a concentration of 400 PFU/mL at 37°C with 5% CO2 for 30 min. Antibody-virus complexes were then added to VAT cells in 12-well plates and incubated for 30 min, then overlaid with 2 mL per well of 0.6% microcrystalline cellulose (Sigma) in minimal essential media (ThermoFisher) supplemented with 0.3% bovine serum albumin (Sigma), and 1% penicillin/streptomycin (ThermoFisher). After 72 hours incubation, plaques were visualized by incubation in 0.25% crystal violet in 100% methanol for 10 minutes. All virus experiments were performed in an approved biosafety level 3 (BSL-3) facility.

### Deep Mutational Scanning

BA.1 full spike deep mutational scanning libraries were designed as described previously^29^. BA.2 full spike deep mutational scanning libraries were designed using the same methods as BA.1 libraries except using BA.2 spike as a template sequence. The sequence of BA.2 spike can be found at https://github.com/dms-vep/SARS-CoV-2_Omicron_BA.2_spike_DMS_COV2-2130/blob/main/library_design/reference_sequences/3332_pH2rU3_ForInd_Omicron_sinobiological_BA2_B11529_Spiked21_T7_CMV_ZsGT2APurR.gb. For antibody escape mapping experiments, each library was incubated for 1 h at 37°C with increasing amounts of COV2-2130 or 2130-1-0114-112 antibodies. For COV2-2130, starting antibody concentration for the BA.1 libraries was 50 µg/ml and increased 4- and 8-fold; for the BA.2 libraries, the starting concentration was 0.32 µg/ml and increased 5- and 25-fold. For 2130-1-0114-112, starting antibody concentration for the BA.1 libraries was 0.16 µg/ml; for the BA.2 library, starting concentration was 0.11 µg/ml and in both cases concentrations were increased 5 and 25-fold. After incubation virus-antibody mix was used to infect HEK-293T-ACE2 cells^56^ and viral genomes were recovered for deep sequencing 12 h after infection. Two biological replicates (virus libraries with independent sets of mutations) were used for each antibody mapping.

Escape for each mutation in the library was quantified using a non-neutralized control as described previously^29^. This analysis uses a biophysical model described previously^63^ and implemented in the polyclonal package found at https://jbloomlab.github.io/polyclonal/. Full analysis pipeline for each antibody can be found at https://dms-vep.github.io/SARS-CoV-2_Omicron_BA.1_spike_DMS_COV2-2130/ for BA.1 libraries and at https://dms-vep.github.io/SARS-CoV-2_Omicron_BA.2_spike_DMS_COV2-2130/ for BA.2 libraries.

### Cryo-EM sample preparation and data collection

The Fab 2130-1-0114-112 and Omicron BA.2 were expressed recombinantly and combined in a molar ration of 1:4 (Ag:Fab). The mixture was incubated overnight at 4°C and purified by gel filtration. 2.2µl of the purified mixture at a concentration of 0.5 mg/mL was applied to glow discharged (30 s at 25mA) grid (300 mesh 1.2/1.3, Quantifoil). The grids were blotted for 3.5 s before plunging into liquid ethane using Vitrobot MK4 (TFS) at 20°C and 100% RH. Grids were screened on a Glacios (TFS) microscope and imaged on Krios operated at 300 keV equipped with a K3 and GIF (Gatan) DED detector using counting mode. Movies were collected at nominal magnification of 130,000X, pixel size of 0.647 Å/pixel and defocus range of 0.8 to 1.8 µm. Grids were exposed at ∼1.09 e^-^/Å^2^/frame resulting in total dose of ∼52.2 e^-^/Å^2^ (**Table EDT2**).

### Cryo-EM data processing

Data processing was performed with Relion 4.0 beta2^64^. Movies were preprocessed with Relion Motioncor2^65^ and CTFFind4^66^. Micrographs with low resolution, high astigmatism, and defocus were removed from the data set. The data set was first manually picked to generate 2D images and then autopicked by Relion template picker^67^ and subject to 2D and 3D classification. Good classes were used for another round of autopicking with Topaz training and Topaz picking^64, 68^. The particles were extracted in a box size of 600 pixel and binned to 96 pixels (pixel size of 4.04 Å/pixel). The particles were subjected to multiple rounds of 2D class averages, 3D initial map and 3D classification without symmetry to obtain a clean homogeneous particle set. This set was re-extracted at a pixel size of 1.516 Å/pixel and was subjected to 3D autorefinement. The data were further re-extracted at a pixel size of 1.29Å/pixel and processed with CTFrefine, polished^69^ and subjected to final 3D autorefinement and postprocessing resulting in ∼3.26Å map. To better resolve the area of interaction between Cov2-RBD/2130-1-0114-112, a focused refinement was performed by particles expansion (C3 symmetry) and signal subtraction with masking around the RBD/2130-1-0114-112. The subtracted particles were subjected to 3D classification without alignment and selected particles were subjected to 3D autorefinement and postprocessing resulting in ∼3.7Å map. Detailed statistics are provided in **Fig. ED7** and **Table EDT2**.

### Model building and refinement

For model building PDB: 7L7E^13^ was used for initial modelling of the RBD and the 2130-1-0114-112 Fv. All models were first docked to the map with Chimera^50^ or ChimeraX^70^. To improve coordinates, the models were subjected to iterative refinement of manual building in Coot^71^ and Phenix^72, 73^. The models were validated with Molprobity^74^ (**Table EDT2**). The EM map and model has been deposited into EMDB (EMD-28198, EMD-28199) and PDB (8EKD).

## Data availability

The EM map and model has been deposited into EMDB (EMD-28198, EMD-28199) and PDB (8EKD). Top antibody sequences are in the accompanying extended data.

## Code availability

Our code is not approved for public release at this time.

## Acknowledgements

This work was performed under the auspices of the U.S. Department of Energy by Lawrence Livermore National Laboratory (LLNL) under contract DE-AC52-07NA27344. The work was supported by the DOD’s Joint Program Executive Office for Chemical, Biological, Radiological and Nuclear Defense, in collaboration with the Defense Health Agency COVID funding initiative under agreement 11647302 in support of the Generative Unconstrained Intelligent Drug Engineering (GUIDE) program, the Defense Advanced Research Projects Agency (DARPA) agreement number HR0011154580 (Amy Jenkins), Laboratory Directed Research and Development programs (20-ERD-032 and 20-ERD-064) at LLNL., and grants and contracts from the NIH (R01 AI157155, NIAID Centers of Excellence for Influenza Research and Response (CEIRR) contract 75N93019C00051). The following reagents were obtained through BEI Resources, NIAID, NIH: SARS-Related Coronavirus 2, Wuhan-Hu-1 Spike D614G-Pseudotyped Lentiviral Kit (NR-53817), and Human Embryonic Kidney Cells (HEK-293T) Expressing Human Angiotensin-Converting Enzyme 2, HEK-293T-hACE2 Cell Line (NR-52511). BD and JDB were supported in part by NIH / NIAID grant R01AI141707. JDB is an Investigator of the Howard Hughes Medical Institute. EM data collections were conducted at the Center for Structural Biology Cryo-EM Facility at Vanderbilt University. The authors wish to thank Amy Jenkins and Jim Brase for significant technical and programmatic contributions and for facilitating this collaborative research effort, Mikel Landajuela and Jiachen Yang for technical feedback, and Elliot Jaffe for editorial contributions. The authors also thank the labs of Prof Sriram Subramaniam and Prof Xinquan Wang for their CryoEM structures of Omicron ahead of release. LLNL-JRNL-839587-DRAFT.

## Contribution Statement

TAD, KTA, ATZ, EYL, FZ, SC, SJZ, EB, CGE, SH, LBT, BWS, AML, SS, MSD, JEC, RHC, and DMF contributed to the conception or design of the study. TAD, KTA, DR, SC, SJZ, EB, SMS, BD, TBE, EC, LSH, LH, DRW, JK-YL, BR, EAS, TW, T-HL, BW, JBC, EAG, BKP, LBT, BWS, JB, MSD, JEC, RHC, and DMF acquired, analyzed, or interpreted data. TAD, ATZ, EYL, FZ, JWG, DV, SN, AL, MSS, RMH, EAG, BKP, and DMF created new software. TAD, KTA, ATZ, EYL, FZ, DR, SC, SJZ, EB, BD, TE, TWB, BKP, BWS, JB, MSD, JEC, RHC, and DMF drafted or substantively revised the manuscript.

## COMPETING FINANCIAL INTERESTS

M.S.D. is a consultant for Inbios, Vir Biotechnology, Ocugen, Moderna and Immunome. The Diamond laboratory has received unrelated funding support in sponsored research agreements from Moderna, Vir Biotechnology, and Emergent BioSolutions. J.E.C. has served as a consultant for Luna Labs USA, Merck Sharp & Dohme Corporation, Emergent Biosolutions, and GlaxoSmithKline, is a member of the Scientific Advisory Board of Meissa Vaccines, a former member of the Scientific Advisory Board of Gigagen (Grifols) and is founder of IDBiologics. The laboratory of J.E.C. received unrelated sponsored research agreements from AstraZeneca, Takeda, and IDBiologics during the conduct of the study. J. D. B. is on the scientific advisory boards of Apriori Bio, Aerium Therapuetics, Invivyd, and the Vaccine Company. Lawrence Livermore National Laboratory, Los Alamos National Laboratory, and Vanderbilt University have applied for patents for some of the antibodies in this paper, for which T.A.D, K.T.A, A.T.Z., E.Y.L., F.Z., A.M.L., R.H.C., J.E.C., and D.M.F. are inventors. Vanderbilt University has licensed certain rights to antibodies in this paper to Astra Zeneca. J. D. B. and B.D. are inventors on Fred Hutch licensed patents related to the deep mutational scanning of viral proteins.

Correspondence and requests for materials should be addressed to D.M.F.

## Extended Data

**Figure ED1:**
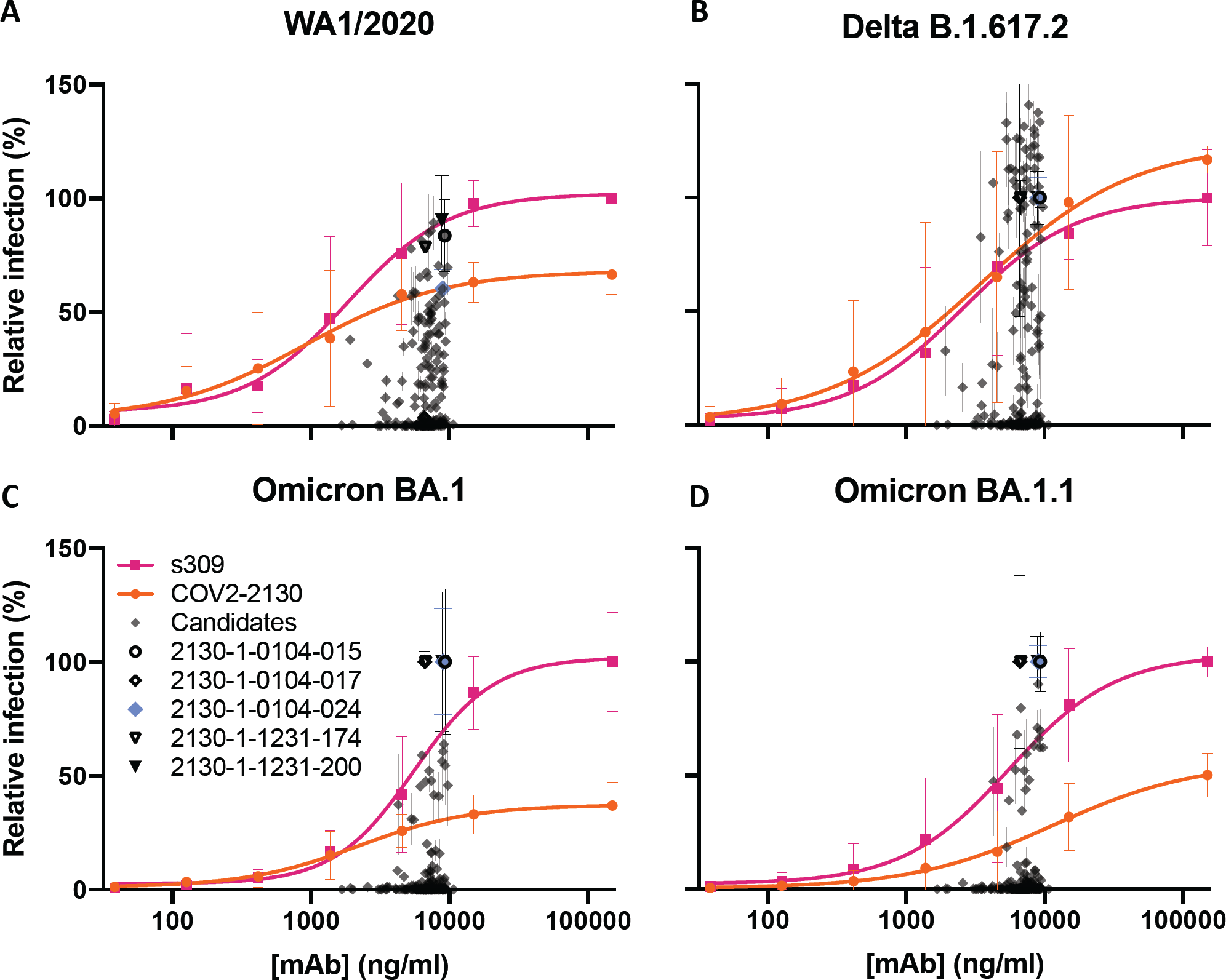
Optimized, single-concentration screening data in Gyrolab immunoassays allow selection of candidates from Set 1 (n=230) for down-stream characterization. Parental mAb COV2-2130 (orange circles) and positive control mAb S309^24^ (magenta squares) serve as references for computationally designed mAbs in single-concentration immunoassays. Target antigens are (A) wild type WA1/2020, (B) Delta, (C) Omicron BA.1 and (D) Omicron BA.1.1. Each screened antibody and antigen combination was evaluated in two replicate assays; each of the controls was replicated in two replicate assays for each of three groups of antibodies, resulting in six replicate assays for each point on the control curves. All error bars indicate standard deviation.

**Figure ED2:**
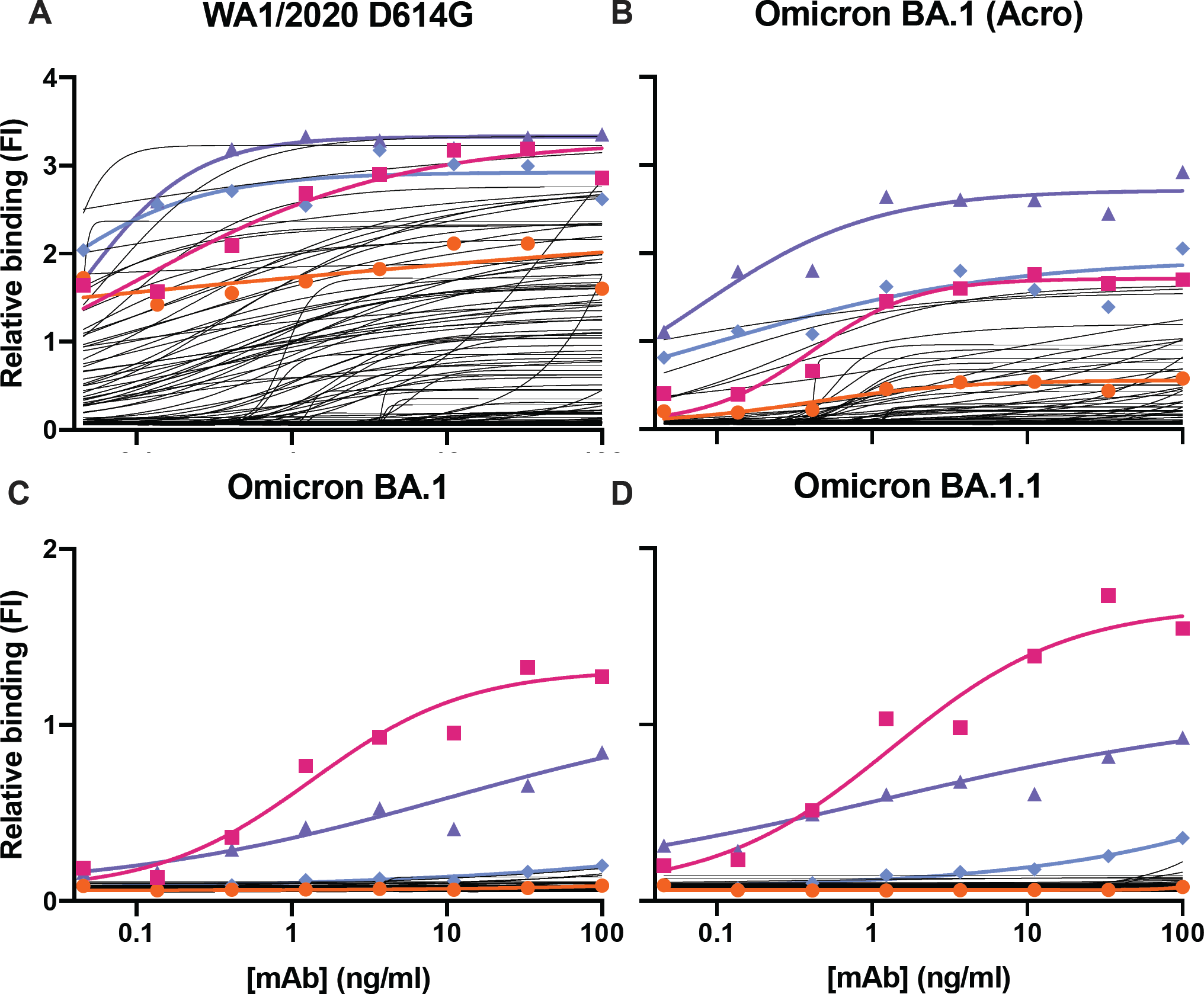
ELISA screening allows set for down selection of candidates from Set 2 (n=204), for down-stream characterization. Parental mAb COV2-2130 (orange circles; 3 technical replicates) and positive control mAb S309 (magenta squares; 3 technical replicates^9^) serve as references for computationally designed mAbs (black curves). Purple triangles are 2130-1-114-112; blue diamonds are 2130-1-0104-024. Each designed antibody had a single measurement (n=1) at each concentration. All curves are 4-parameter logistic fits, produced in GraphPad Prism. Target antigens are RBD from (A) wild type WA1/2020, (B) Omicron BA.1 (Acro), which is biotinylated, (C) Omicron BA.1, and (D) Omicron BA.1.1. For the biotinylated antigen in (B)(Acro Biosystems, cat. SPD-C82E4), an additional coat and wash cycle was required to prepare the ELISA plate with streptavidin.

**Table EDT1:**
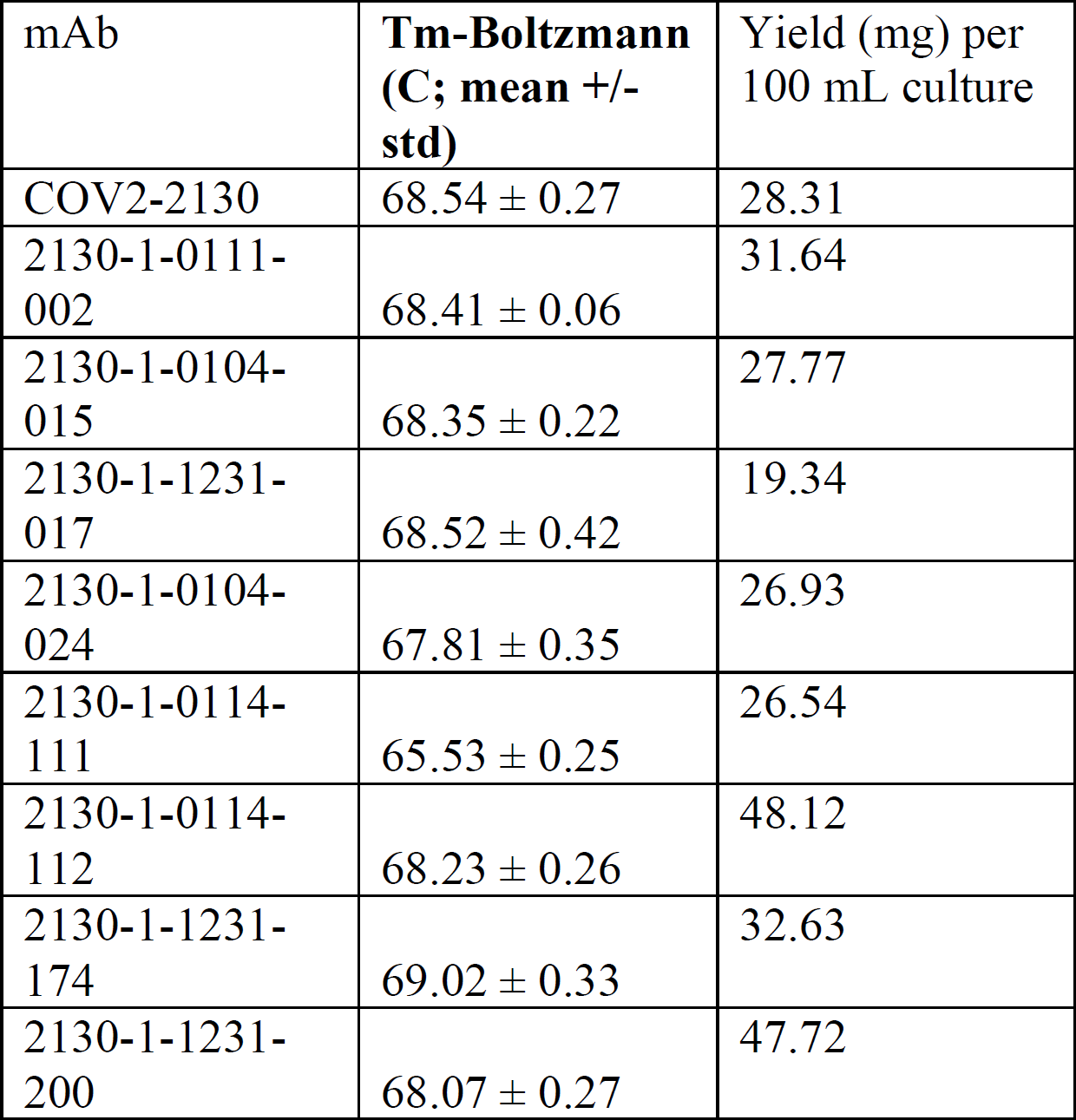
Thermal stability and expression yield of selected IgGs. Melting temperature (Tm) was determined using a fluorescence-based protein thermal shift assay (GloMelt^TM^, Biotium). Yield was determined by measuring optical density at 280nm and deriving antibody quantity using the calculated extinction coefficient.

**Figure ED3:**
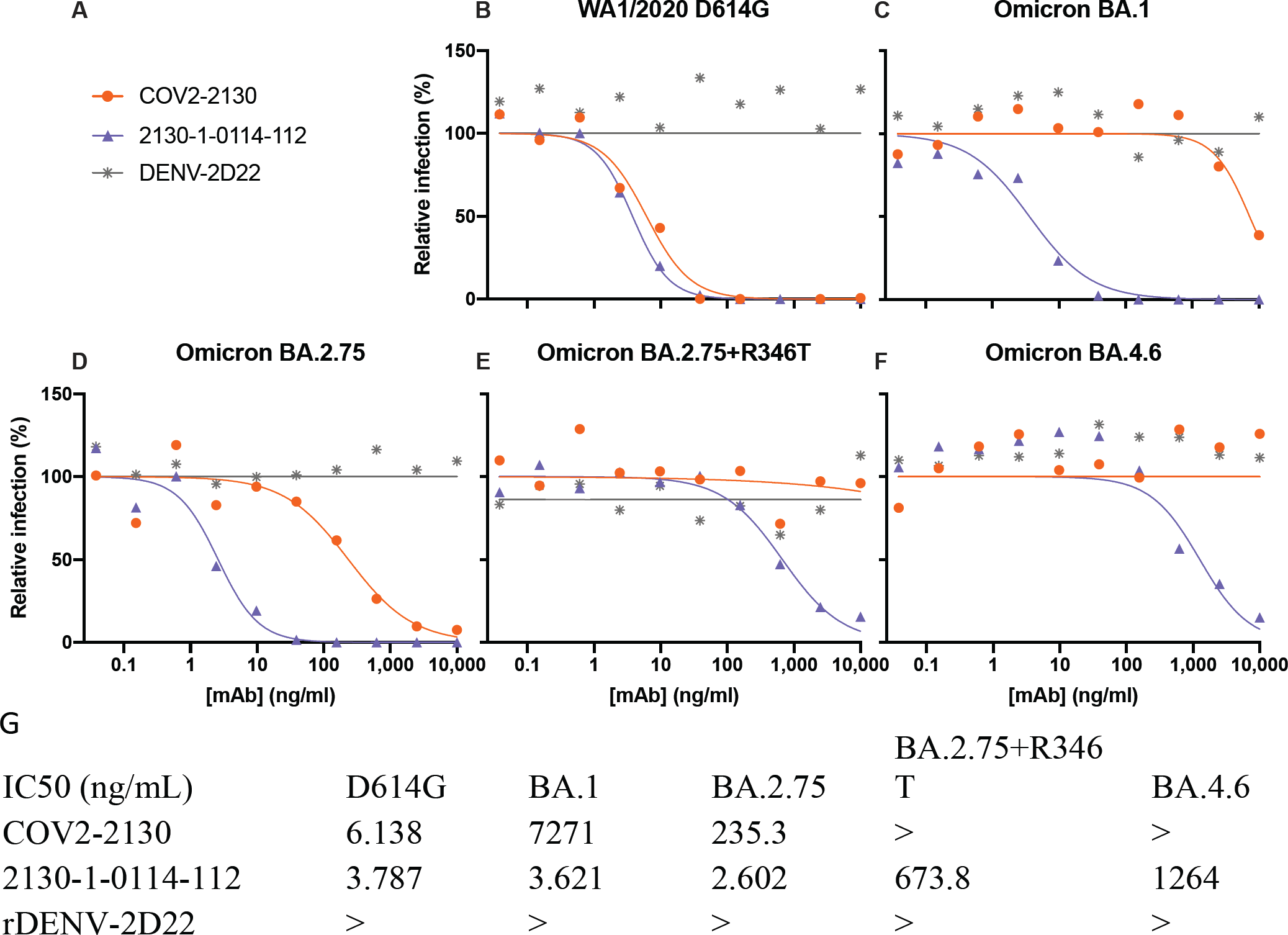
Normalized pseudoviral neutralization of recently emerged VOCs by 2130-1-0114-112. (B) COV2-2130 and 2130-1-0114-112 potently neutralize WA1/2020 D614G and (C) only 2130-1-0114-112 potently neutralizes BA.1, consistent with other pseudoviral neutralization assays. (D) 2130-1-0114-112 potently neutralizes BA.2.75, outperforming COV2-2130 by 90-fold. (E-F) 2130-1-0114-112 loses substantial potency in the context of BA.4.6 and artificially-produced BA.2.75+R346T but retains measurable neutralization, demonstrating mitigation of this critical weakness of COV2-2130. All sub-figures (B-F) show the mean of two technical replicates and curves show a fitted four-parameter logistic curve with fixed minimum and maximum values. (G) Neutralization IC50 values in ng/ml. “>” indicates IC50 greater than 10,000 ng/mL. All analysis conducted in GraphPad Prism.

**Figure ED4.**
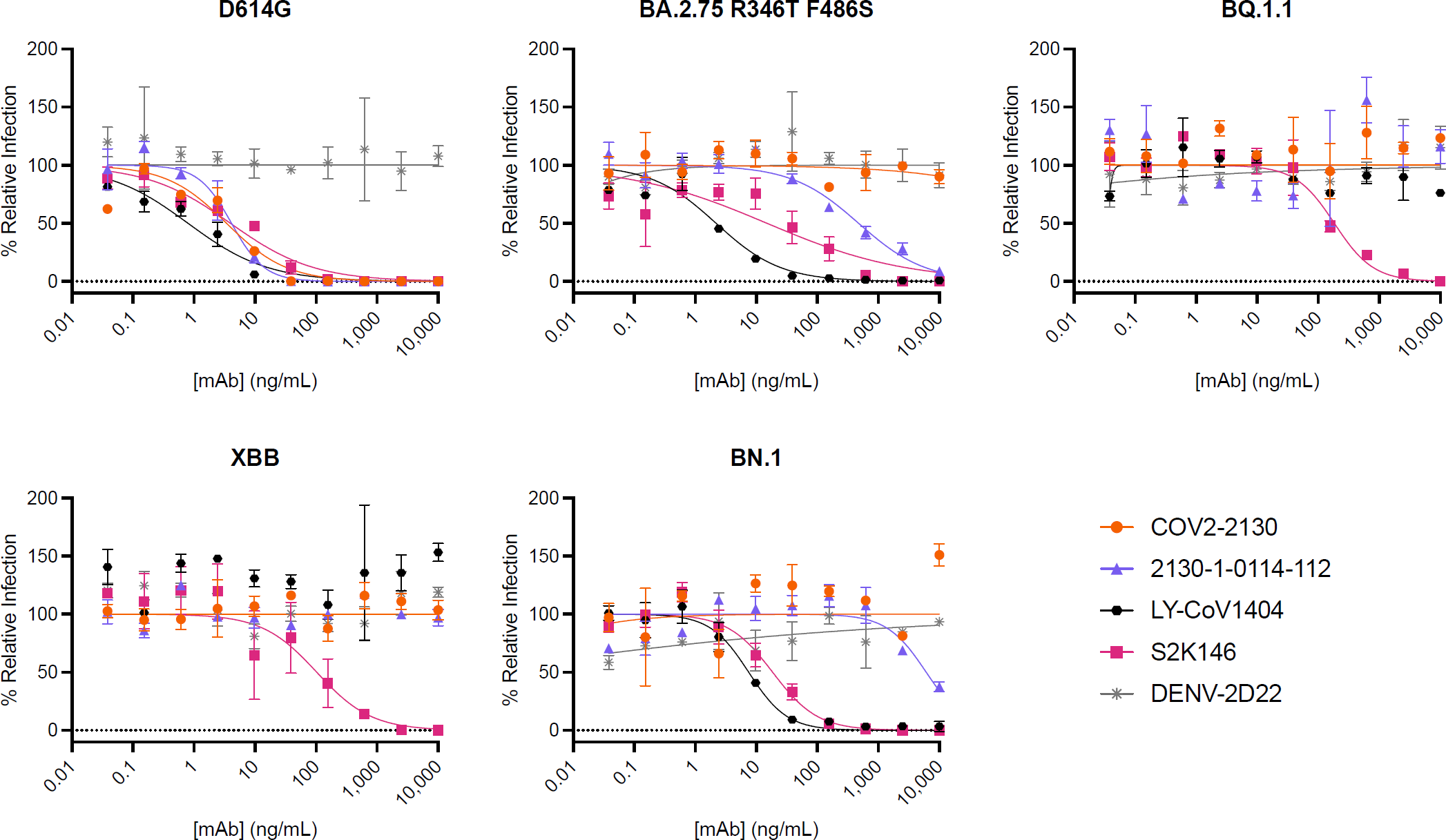
Normalized pseudoviral neutralization of recently emerged VOCs. Pseudoviral neutralization assays testing COV2-2130 and 2130-1-0114-112 against D614G and BA.2.75 R346T F486S, BQ.1.1, XBB, and BN.1 variants. COV2-2130 has no detectable activity against all variants tested. In contrast, 2130-1-0114-112 maintains neutralization of BA.2.75 R346T F486S, but loses detectable neutralization activity against BQ.1.1 and XBB and exhibits a near-complete loss of neutralization activity against BN.1. All data points represent the mean of two technical replicates, while error bars denote the standard deviation. Curves represent a four-parameter logistic curve fit to the data using GraphPad Prism with fixed minimum and maximum values (0 and 100, respectively).

**Figure ED5:**
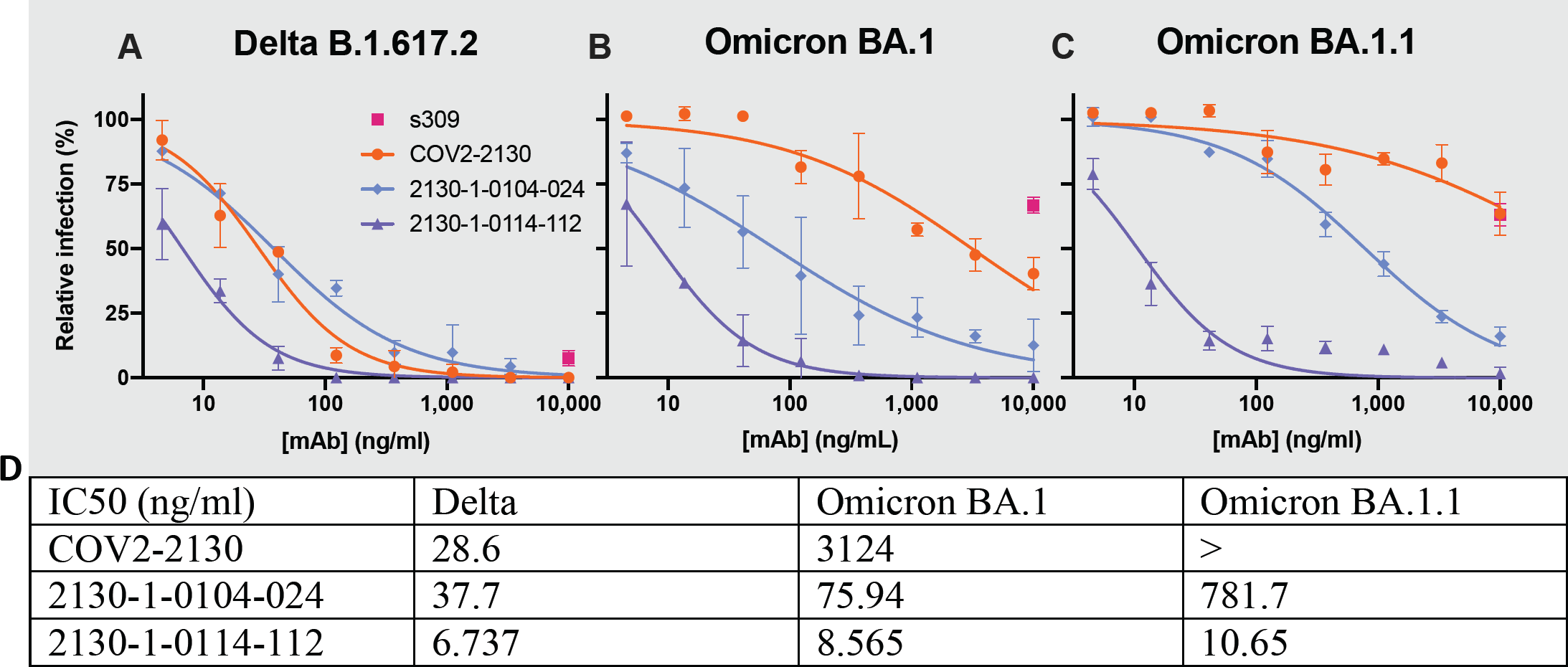
Authentic virus neutralization in plaque assays using Vero E6-TMPRSS2-T2A-ACE2 (VAT) cells. Plaque assay neutralization of (A) Delta, (B) Omicron BA.1 and (C) Omicron BA.1.1 viruses. Data are represented as the normalized infection of mAb-treated virus to virus treated with control human IgG (Invitrogen). For S309, each point shows the mean of four technical replicates; all other points are means of two technical replicates; error bars show standard deviation. Curves are two-parameter (IC50, hill-slope) logistic fits to normalized response. (D) IC50 values (ng/ml, “>” indicates IC50 greater than 10,000 ng/mL) show that 2130-1-0104-024, while having only two mutations from COV2-2130, remains potent against BA.1 and suffers a 20-times loss in potency against BA.1.1. 2130-1-0114-112 is strongly potent against all three tested variants. All analysis performed in GraphPad Prism.

**Figure ED6:**
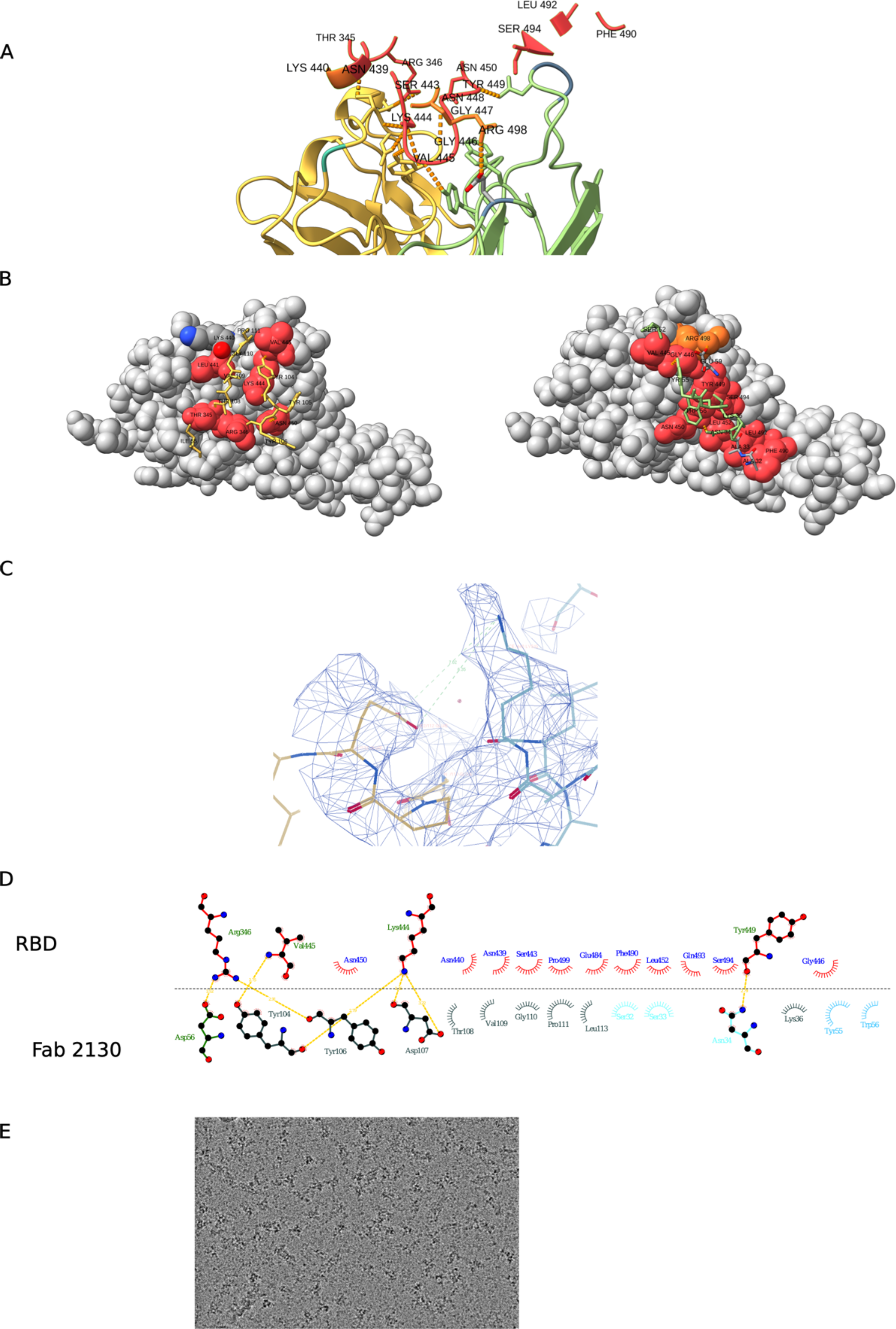
Cryo-EM structure of neutralizing antibody 2130-1-0114-112 in complex with Omicron BA.2 RBD. (A) RBD residues in 7 Å distance form 2130-1-0114-112. RBD in red, with BA.2 mutated residues in orange, and 2130-1-0114-112 in yellow/green. (B) Left, 3D representation of the interaction plot between RBD and 2130-1-0114-112 HC. 2130-1-0114-112 is shown as stick (yellow) and RBD as gray spheres with the contact residues in red. Contact residues labelled and numbered. Right, 3D representation of the interaction plot between RBD and 2130-1-0114-112 LC. 2130-1-0114-112 show as stick (green) and RBD as gray sphere with the contact residues in red. Contact residues are labelled and numbered. (C) Glu112 and Lys440 with the EM map and distance between the sidechains. (D) Fab COV-2130 paratope and epitope residues involved in hydrogen bonding (dashed lines) and hydrophobic interactions. Hydrophobic interactions residues are shown as curved lines with rays. Atoms shown as circles, with oxygen red, carbon black, and nitrogen blue. Image was created with Ligplot+. (E) Representative micrograph.

**Figure ED7:**
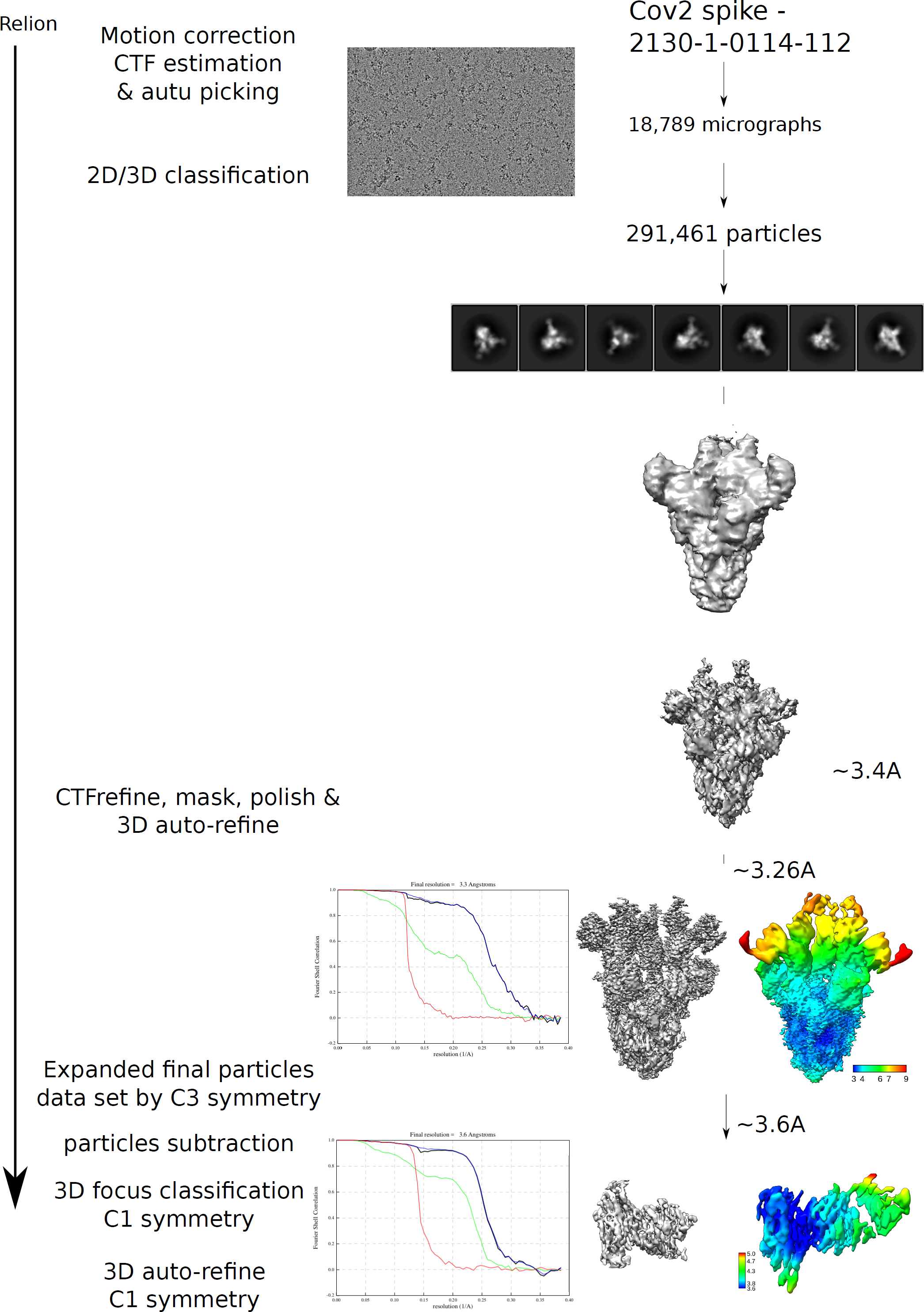
CryoEM workflow of SARS-CoV-2 BA.2 spike bound to Fab. At the bottom, Gold-standard Fourier shell correlation curves and maps colored by local resolution calculated using Relion, before and after local refinement.

**Table EDT2:**
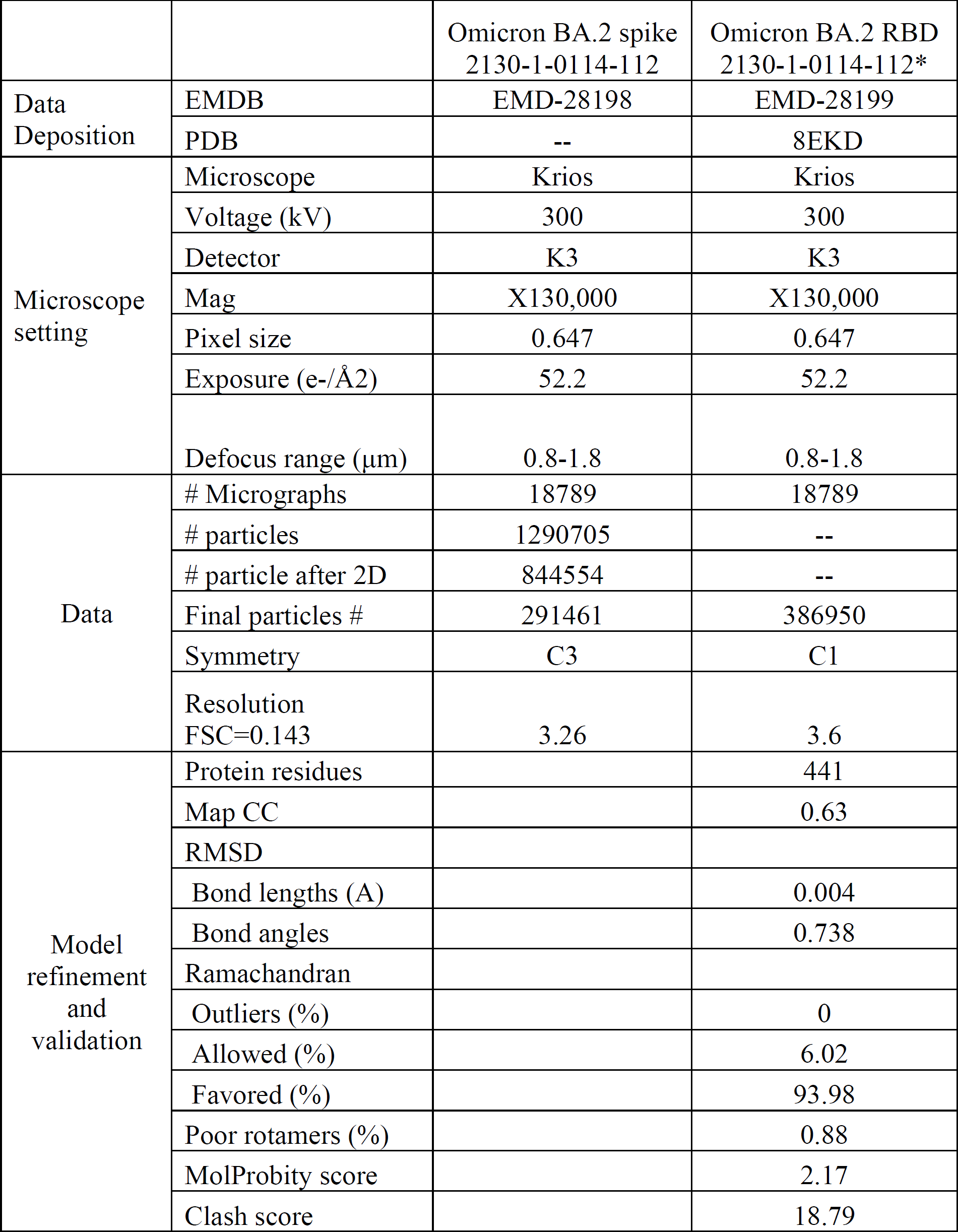
Parameters of CryoEM procedures. *: Focus refinement

**Figure ED8:**
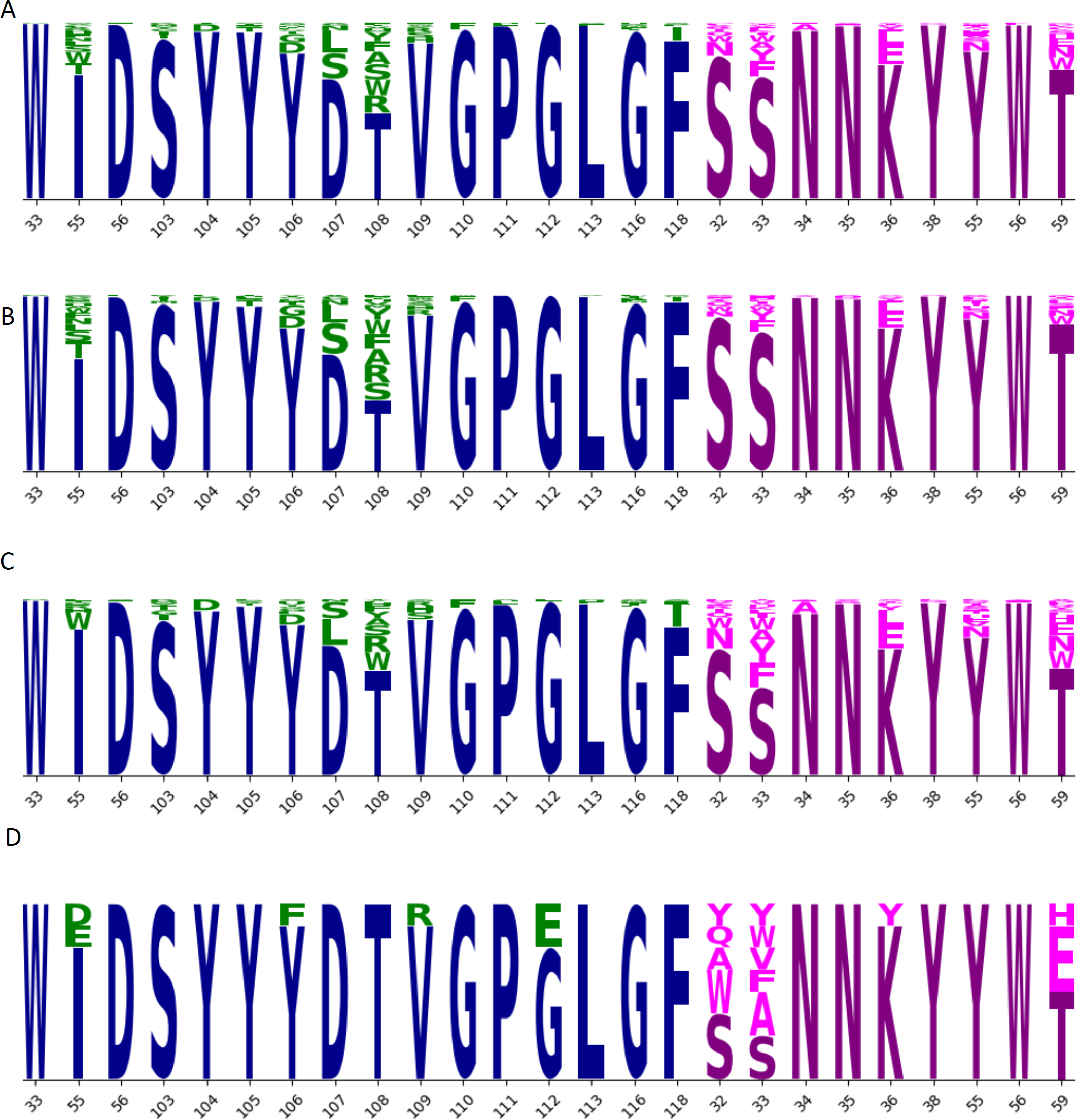
Sequence logos. The set of 376 designed IgG (A) includes mutation at 16 positions in the heavy chain (blue; mutations in green) and 9 positions in the light chain (magenta; mutations in pink). Height of each letter is proportional to the frequency of the amino acid in the group. This set of 376 sequences is divided into two overlapping sets, Set 1 (B; n=230) and Set 2 (C; n=204). From these two sets, a set of eight sequences (D) was selected for further evaluation. Selected sequences show reduction in mutations throughout the CDRH3 residues (103-118) mutated in (A), especially in residues 103-108. After eliminating sequences 2130-1-1231-194 (IH55W) and 2130-1-1231-199 (SL33Y), we produced the remaining eight sequences at larger scale and evaluated their thermostability and binding performance (Fig 2).

**Fig. ED9:**
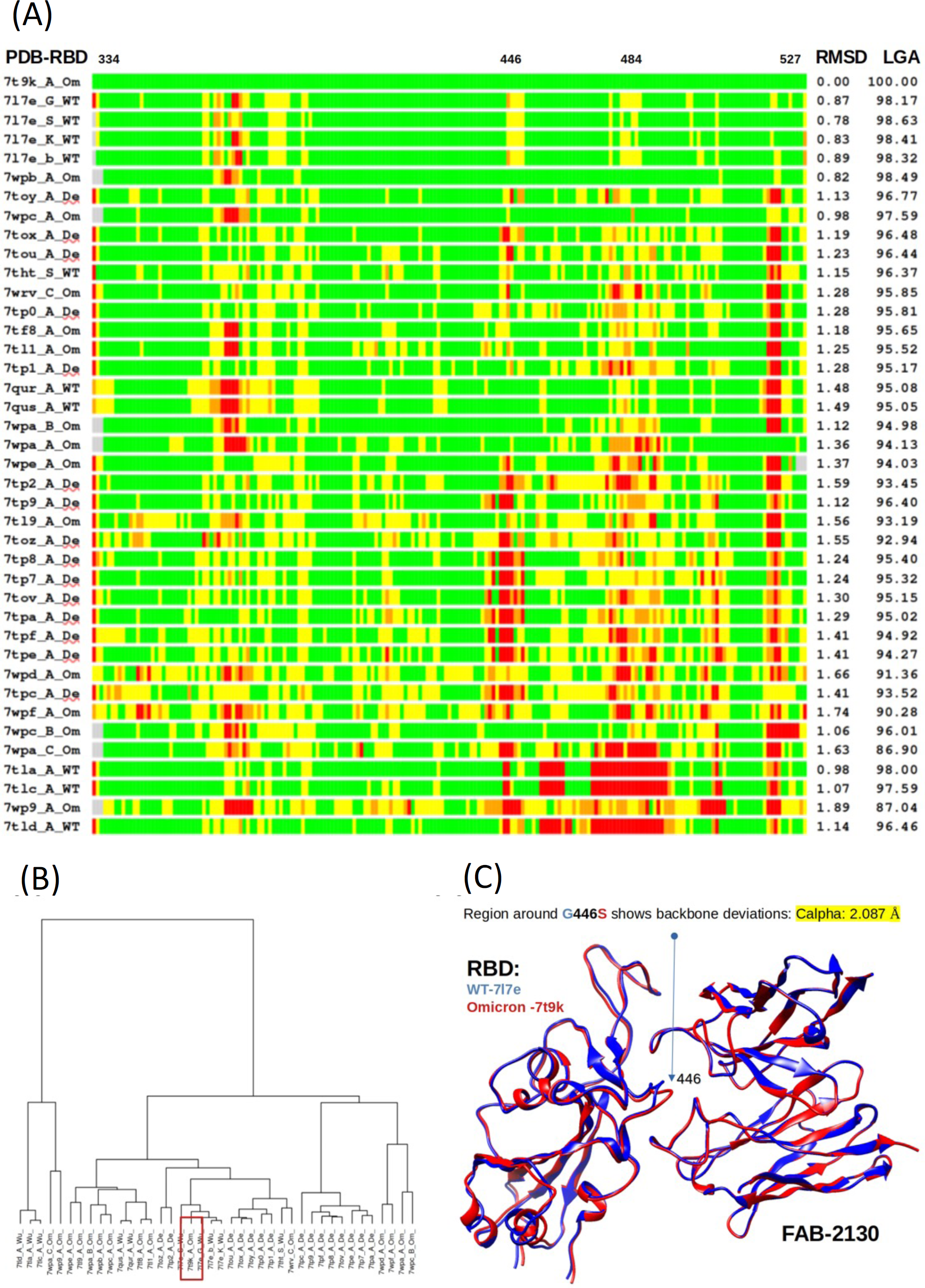
Bar plot representation of superpositions of the reference Omicron RBD structure (PDB ID 7t9k chain A) with 40 RBDs from WT, Omicron or Delta variants and comparison of experimentally solved RBD structures. (A) The RMSD (Å) and LGA structure similarity scores (0-100) were calculated against the reference structure are provided in the right columns. Deviations of < 1Å are shown in green, 1-2 Å in yellow, 2-4 Å in orange, and > 4Å in red. The regions in RBD within the RBD-Fab interface where the major structural deviations between Omicron, Delta and WT are observed (positions 446 and 484) are marked at the top. (B) Structural clustering by LGA of 40 experimentally-solved RBDs. A red rectangle marks an identified centroid (an RBD from Omicron PDB ID 7t9k chain A) (C) A structure of WT RBD-COV2-2130 (PDB ID 7L7E, blue) superimposed with a model of Omicron RBD-COV2-2130 (Derived from PDB ID 7T9K, red). A significant deviation between two models is observed in the RBD-Fab interface in the region surrounding a mutation position WT to Omicron G446S (arrow).

**Figure ED10.**
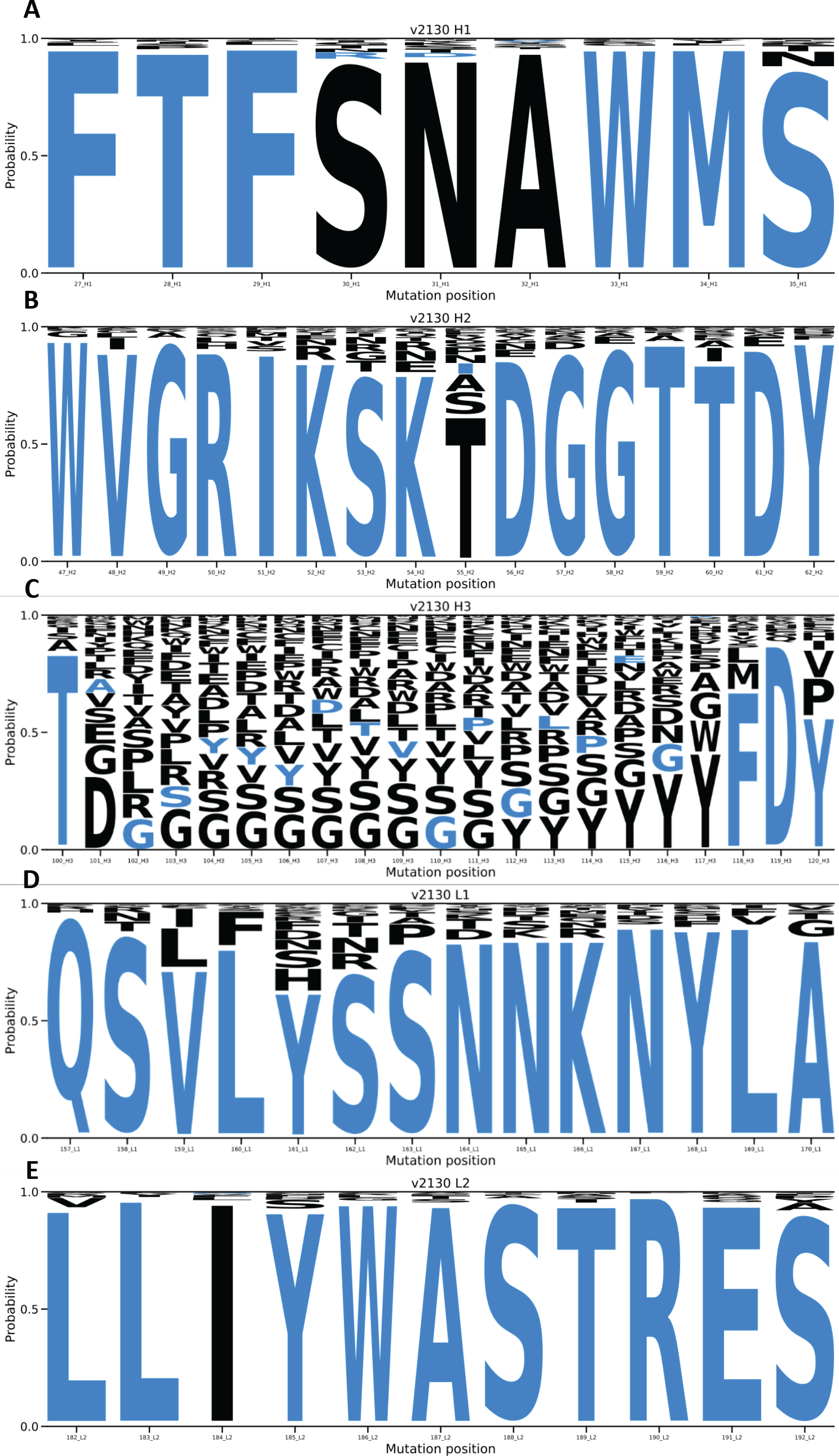
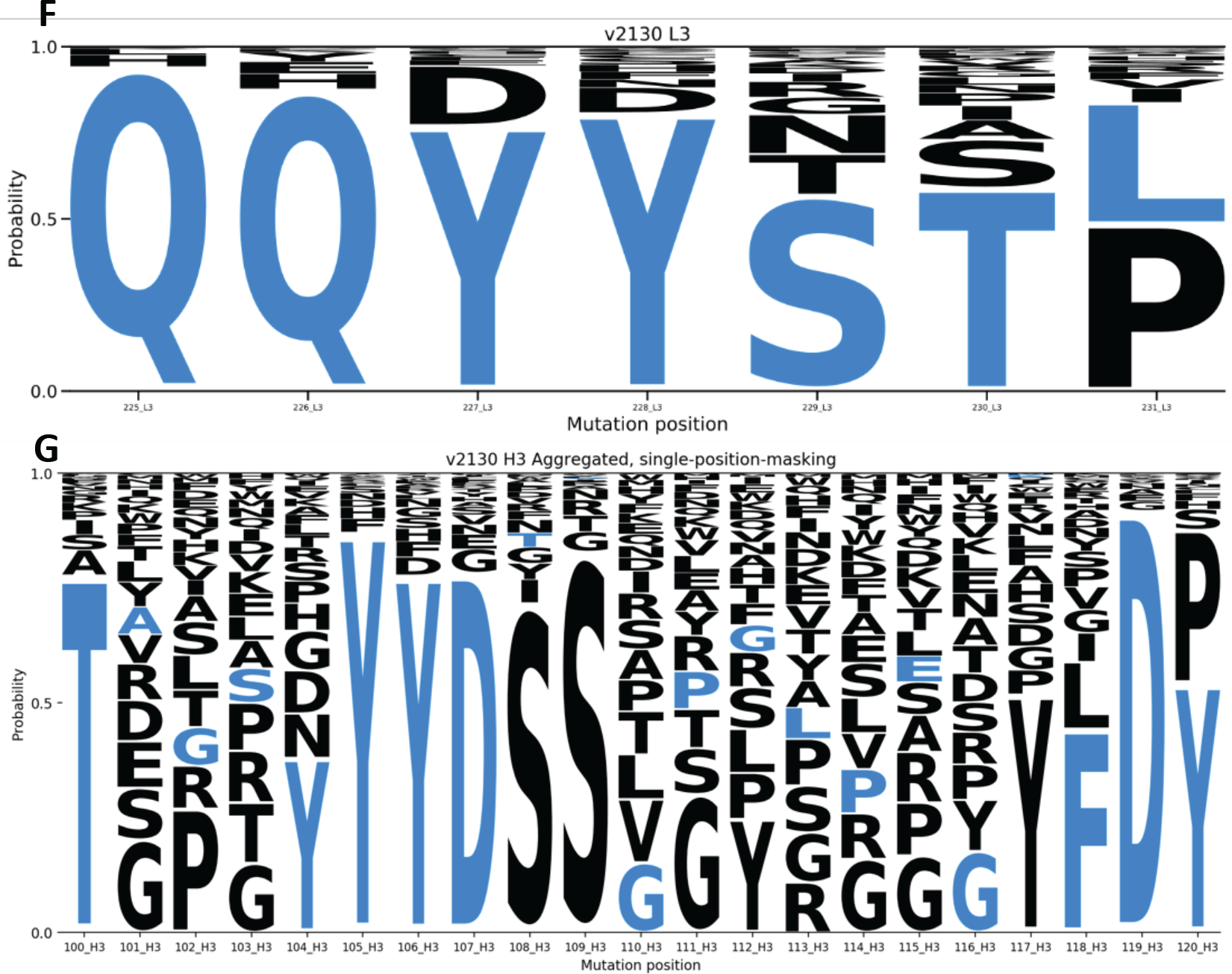
The AbBERT model can be used to generate whole CDR replacements in *COV2-2130*. For each sub-figure, 1,000 sequences were sampled by masking and replacing residues in loop regions: (A) CDRH1, (B) CDRH2, (C) CDRH3, (D) CDRL1, (E) CDRL2, and (F) CDRL3. Wild type COV2-2130 residues are in blue; mutations are in black. Letter heights correspond to frequency in the 1,000 sampled sequences. Because the residues are numbered in concatenated order, all residue numbers in the light chain are offset from their conventional values by +130, the length of the heavy chain. (G) Masking and replacing individual CDRH3 residues, executed separately for each position shown here, preserves considerably more structure and information about the COV2-2130 CDRH3 than complete masking and reconstitution of the entire CDRH3 as in (C).

Selected Sequence Records (Corresponding to Fig. ED8D)

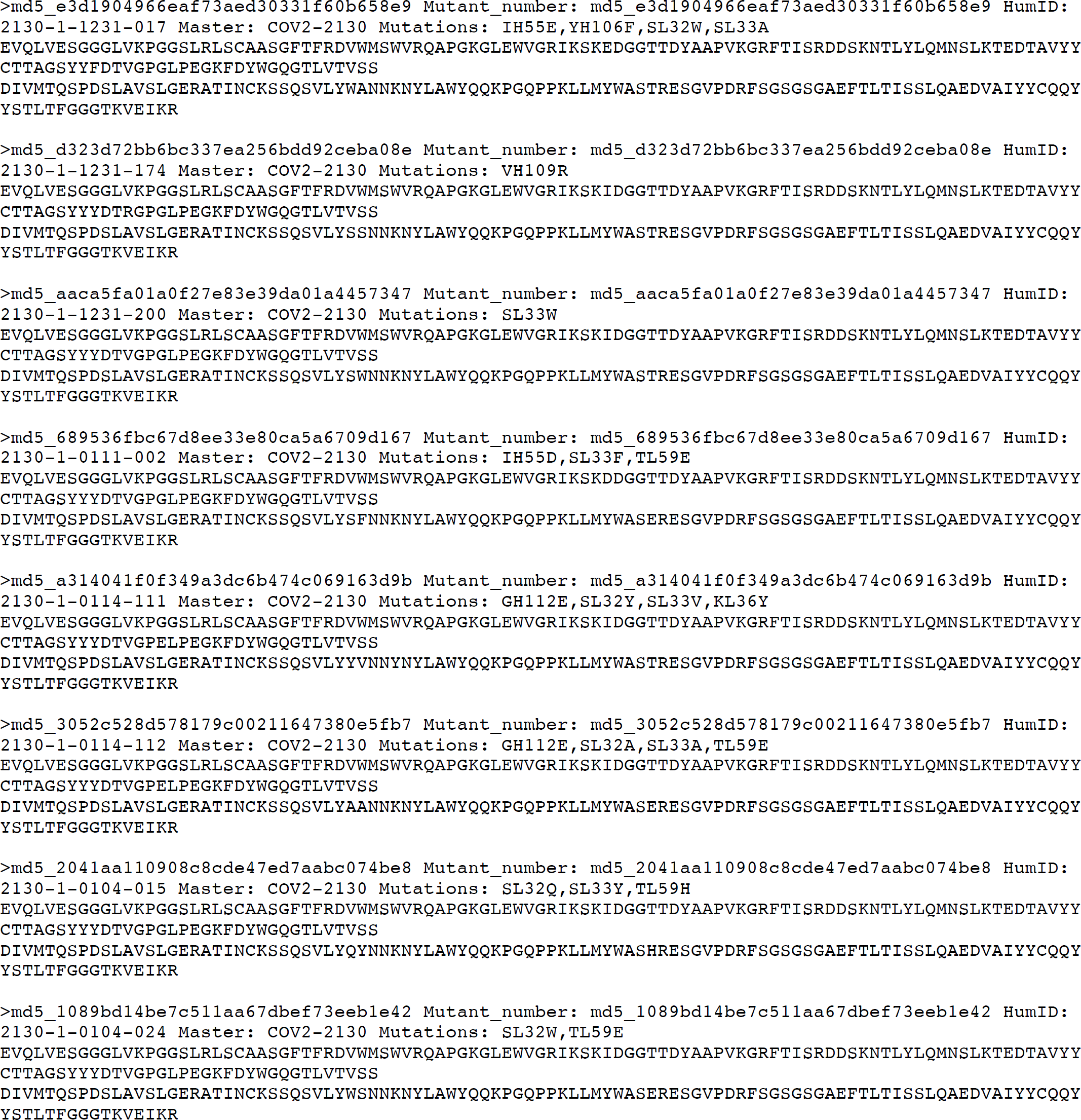

